# Distinct modes of sequence evolution and epigenetic modifications underpin the origins of *Starship*-mediated variation in *Pyricularia* fungal plant pathogens

**DOI:** 10.64898/2026.01.28.702382

**Authors:** Samuel O’Donnell, Aidan McVey, Barbara Valent, Sanzhen Liu, Emile Gluck-Thaler, David E. Cook

**Author notes:** Authors contributed equally (Samuel O’Donnell, Aidan McVey).

## Abstract

Fungal pathogens display remarkable variation in genome content and organization that directly impacts their survival and host interactions. Although numerous models have been proposed to explain the origins of this variation, they generally fail to explain or predict the mechanisms that generate the genome variation observed in natural populations. *Starships* are a recently discovered group of giant fungal transposons that carry dozens of genes as cargo and horizontally transfer both within and between species. Here, we identify the features of a newly defined “*Starship* compartment” in the major fungal plant pathogen *Pyricularia oryzae.* We test the hypothesis that the *Starship* compartment makes distinct contributions to fungal genome evolution by explicitly comparing its transferability, mutability, and epigenetic modifications with those of the canonical core and accessory compartments. To enable this, we developed an updated and user-friendly version of the annotation tool stargraph for the comprehensive annotation of *Starships* and *Starship*-like regions. Using this approach, we identified two distinct families of *Starships* and related *Starship*-like regions in *P. oryzae* that differ in their activity, impacts on genome organization, modes of sequence evolution, and epigenetic modifications. Elements from the more active family exhibit higher rates of structural variation than all other genomic compartments in the predominantly clonal isolates infecting rice. Both families of *Starships* encode specific suites of known effector sequences that contribute to plant disease and *Starship* activity accounts for avirulence gene turnover, which suggests that evolutionary change within the *Starship* compartment may subsequently impact the evolution of plant-fungal interactions. *Starships* from the more active family have repeatedly transferred across the *Pyricularia* genus and tend to be depleted in heterochromatic histone modifications and repeat-induced point mutations. However, contrasting histone modification profiles in this family suggests a genomic conflict between silencing or maintaining *Starship* activity. Our findings demonstrate that variation in the mode of sequence diversification and epigenetic modification within the *Starship* compartment underpins the impacts of these giant transposons on fungal genome evolution. We argue for the explicit consideration of not only the *Starship* compartment but of element-specific dynamics when investigating the evolution of host-fungal interactions.

## INTRODUCTION

Fungal pathogens display remarkable variation in genome content and organization, owing to their fast generation times, large numbers of offspring, and diverse modes of DNA exchange and inheritance. Yet despite the ubiquity of such variation in fungal genomes, basic facts about its origins and consequences remain unknown. A variety of models and terms have consequently been proposed to describe how genomic variation originates, how it is organized, and how it impacts the evolution of fungal pathogenesis. One example is the designation of core and accessory genes that respectively make up core and accessory genomes, which are defined by their degree of presence/absence variation (PAV) observed at the population level (McNally et al. 2016). Another model prevalent in literature of fungal and oomycete eukaryotic pathogens describes a 2-speed genome model composed of genes that experience “fast” and “slow” rates of evolution (Raffaele et al. 2010b; Dong et al. 2015). The two-speed genome model describes how individual genes in gene dense regions with few repetitive elements (i.e., genes in close linear proximity to other genes) experience slower rates of sequence change while genes in gene sparse regions near repetitive elements show characteristics of elevated diversity such as higher PAV and elevated rates of sequence variation (Raffaele et al. 2010a). Although conceptually powerful, several fungal pathogens do not display a 2-speed genome organization and the fast and slow speeds are likely part of a multi-speed spectrum (Sánchez-Vallet et al. 2018; Frantzeskakis et al. 2019; Torres et al. 2020). A limitation of the core and accessory and two-speed genome models is that they do not provide mechanisms or predictions for how genome variation is created or maintained.

Alternatively, the concept of genome compartmentalization is a simple extension of the core/accessory and slow/fast genome models that refines our ability to understand and predict the evolution of genomic variation (Frantzeskakis et al. 2019). In the compartmentalization model, core compartments are physically contiguous regions of DNA with low PAV that are characterized by relatively low densities of repetitive elements and a high density of conserved genes responsible for basic cellular and biological functions. In contrast, accessory compartments are regions with high PAV, characterized by higher repetitive element content, lower gene content, and genes associated with adaptability and host interactions. Genome compartmentalization is not strictly a binary model, and the core and accessory compartments can be thought to represent opposite ends of a spectrum defined by variation across multiple genomic features (e.g. repeat content, PAV, mutation rate, etc). The genome compartmentalization model offers a way to quantify genomic variation within a contiguous DNA region and investigate the underlying mechanism of creation or maintenance. For example, the activity and subsequent regulation and silencing of transposable elements is linked to the development of genomes with compartmentalized organization across many eukaryotes (Gozashti et al. 2024).

An example of the utility of the genome compartmentalization model is the ability to incorporate epigenetic information into models of genome evolution (Fouché et al. 2022). Indeed, joint investigations of epigenetic and genetic variation have revealed intriguing associations between these dimensions of genomic diversity (Connolly et al. 2013; Schotanus et al. 2015; Wang et al. 2017; Cook et al. 2020). For example, selection for fine-tuned epigenetic regulation of effector genes (coding sequences mediating host infection) and secondary metabolite gene clusters in fungal pathogen genomes has been proposed to result in the co-localization of these loci with epigenetically silenced transposable elements (Fouché et al. 2022). These effector- and TE-rich accessory compartments display specific patterns of histone modifications, such as trimethylation of H3-Lys27 (H3K27me3), H4K20me3, Ash1-mediated H3K36me3, and H3K9me3 (Soyer et al. 2014; Fokkens et al. 2018; Rowe et al. 2023; Möller et al. 2023; Xu et al. 2024). These four histone marks are largely responsible for the formation of heterochromatin, which are DNA regions that are constitutively repressed (e.g., lack transcription and DNA accessibility) or regions that are facultatively repressed allowing for condition-specific DNA activity (Chujo and Scott 2014; Wang et al. 2020; Zhang et al. 2021; Kramer et al. 2023). While the impact of heterochromatin on transcription is well documented, there is increasing evidence that heterochromatin is also associated with variation in the rate of sequence evolution, a key feature of many models of genome evolution and compartmentalization. For example, mutation accumulation experiments show that DNA marked by heterochromatic histone modifications experience higher rates of mutation accumulation (Habig et al. 2021; Peña et al. 2023). In *Pyricularia oryzae* (syn. *Magnaporthe oryzae*), the causative agent of blast disease on monocots, the rate of synonymous substitutions in genes located in H3K27me3 marked regions is significantly higher than genes not in H3K27me3 marked regions for multiple strains (Rowe et al. 2023). Genetic perturbation affecting heterochromatic marks lead to more and less frequent loss of specific accessory chromosomes in the fungal pathogen *Zymoseptoria tritici*, further suggesting the epigenome plays a role in sequence maintenance (Möller et al. 2019). Thus, multiple lines of evidence support the conclusion that particular types of epigenetic modifications are associated with increased genome variability and particular genomic compartments in fungi. What remains unclear is the degree of cause and effect; that is, how variation in the epigenome directly mediates rates of genome evolution and compartmentalization.

Despite its improvement over the core/accessory and fast/slow genome models, a limitation of the compartmentalization model is that it implies all accessory regions have a similar origin and impact on the genome. One avenue for enriching these models comes from the recent discovery of a previously unknown family of giant transposons called *Starships* (Urquhart et al. 2022; Vogan et al. 2021). These unusual elements are widespread in the *Pezizomycotina* subphylum of filamentous fungi, which contains the vast majority of human and animal fungal pathogens (Gluck-Thaler and Vogan 2024). *Starships* do not easily fit into any existing model of fungal genome evolution because unlike other physically contiguous genomic regions, they are mobile DNA capable not only of transposition but of horizontally transferring themselves across species (A. S. Urquhart et al. 2024; Bucknell et al. 2024; O’Donnell et al. 2025b), including under lab conditions between fungal genera separated by >100 million years of evolution (Urquhart et al. 2025). Yet *Starships* do not neatly fit into the category of repetitive elements either because unlike smaller transposons, they consist of much more than the genes necessary for their mobility. *Starships* often span hundreds of kilobases and carry dozens of protein-coding genes with notable and predicted phenotypes important for fungal adaptation, such as secondary metabolite biosynthesis (Gluck-Thaler et al. 2025), formaldehyde resistance, heavy metal tolerance (Urquhart et al. 2022; A. S. Urquhart et al. 2024), spore killing gene drives (Vogan et al. 2021), lactose metabolism (O’Donnell et al. 2025a), anti-fungal drug resistance (O’Donnell et al. 2025a), and host-specific pathogenicity (Gourlie et al. 2022; Sato et al. 2025). The possibility for active transposition and horizontal transfer coupled with cargo-carrying abilities prompted a recent proposal that *Starships* should be considered a separate category of genomic compartment alongside the core and accessory since these traits set them apart from other genomic regions while providing a distinct mechanism for generating genomic variation (A. Urquhart et al. 2024). We propose that incorporating *Starships* into models of genome organization and evolution will improve explanation of extant variation and prediction of future genomic changes Here, we investigate the genetic, epigenetic, and evolutionary features of the *Starship* compartment through explicit comparisons with the core and accessory compartments. Our overarching goal is to understand how the *Starship* compartment impacts broader trends in fungal genome organization and the evolution of plant-fungal interactions. We used the *P. oryzae* pathosystem as a model because it is well suited to study the impact of *Starships* on fungal evolution. First, there are diverse host adapted pathotypes of *P. oryzae* that threaten global food production, including isolates that infect rice (*Oryza*, PoO), wheat (*Triticum*, PoT), ryegrass (*Lolium*, PoL), and finger millet (*Eleusine*, PoE) (Gladieux, Condon, et al. 2018; Valent 2021). The rice infecting isolates spread worldwide as clonal lineages and have relatively low sequence diversity (Zhong et al. 2018; Latorre et al. 2020), while *Lolium* and especially wheat infecting isolates, have emerged more recently and contain much higher levels of genetic diversity (Gladieux, Condon, et al. 2018; Rahnama et al. 2023). *P. oryzae* also has many examples of effector PAV and sequence diversification (Yoshida et al. 2009; 2016; Latorre et al. 2020; Le Naour—Vernet et al. 2023), transposable element linked diversification (Yoshida et al. 2016), effector translocation events (Latorre et al. 2020), and diverse mini-chromosomes that mediate horizontal gene transfer (Peng et al. 2019; Langner et al. 2021). Lastly, there is evidence that *P. oryzae* strains exchange genes and DNA fragments with other related species (Chuma et al. 2011; Kobayashi et al. 2023). Thus, *P. oryzae*’s genome variation coupled with variation in its host-pathogen interactions makes it an excellent model system to evaluate the utility of expanding the concept of genome compartmentalization to include *Starships*.

To overcome existing limitations in *Starship* annotation (see below), we developed an updated and user-friendly version of the “stargraph” annotation tool that we used to more expansively identify *Starships* and *Starship*-like regions (SLRs), which together make up the *Starship* compartment, in 13 *P. oryzae* complete genome assemblies. We find that activity and variation within the *Starship* compartment shapes extant patterns of genome diversity across four major host infecting lineages of *P. oryzae*. We identified two families of *Starships* and related SLRs with markedly different mobility, sequence diversification, and histone modification profiles among strains. The *Starship* family with more frequent horizontal transfer events also has the highest rate of large indel accumulation and sequence diversification compared to all other genomic compartments, suggesting that elements from this family are hotspots for genomic variation. This highlights the diversity-generating potential of the *Starship* compartment, especially in asexual lineages. The most active *Starship* family also displays significant variation in epigenetic marks, suggesting a genomic conflict between maintaining genome integrity and sequence diversification within *P. oryzae*. Our results suggest that the *Starship* compartment is a driving force in fungal genome evolution and organization, creating variable regions, variable genome content, and altered epigenomes contributing to strain heterogeneity with potential to impact host-pathogen interactions.

## RESULTS

### The *Starship* compartment accounts for upwards of 5% of the *P. oryzae* genome

To understand the impact of *Starships* on the evolution of genome variation in this major plant pathogen, we analyzed 13 *P. oryzae* long-read assembled genomes representing strains from four host adapted pathotypes (Table S1). Two complementary pipelines for *Starship* annotation were used (Gluck-Thaler and Vogan 2024) (Fig. 1A). The first, called “starfish”, leverages the fact that *Starships* encode a tyrosine recombinase (*tyrR*) that is necessary for the transposition of the entire element (Urquhart et al. 2023), and identifies candidate *Starships* based on *tyrR* presence at the boundary of a large presence/absence variant (PAV) region from pairwise whole genome alignments (Gluck-Thaler and Vogan 2024). The terminal *tyrR* at the boundary of the element is referred to as the captain to distinguish it from other *tyrR* sequences that may be present in the element. We observed through manual inspections that starfish’s strict requirements for a *tyrR* at one of the boundaries of the PAV and for a clean insertion flanked by alignable regions identified high confidence *Starships*, but the approach did not capture all PAVs with predicted *tyrR* sequences. Therefore, to more precisely quantify the impact of *Starships* on genome evolution, we further developed a previously described and more permissive methodology for the identification of *Starships* into a user-friendly pipeline called “stargraph” (O’Donnell et al. 2025a) (see methods; Fig. S1). Stargraph parses all vs. all whole genome alignments stored in the pangenome variation graph format to identify both *Starships* and *Starship*-like regions (SLRs), the latter of which we define as PAV regions that contain *tyrRs* and are thus likely *Starship*-derived but do not necessarily have well-defined insertion sites. We did not assign any *tyrRs* in SLRs a “captain” status, as this designation is reserved for *tyrRs* at the 5’ boundary of an element with a defined insertion site.

**Figure 1.**
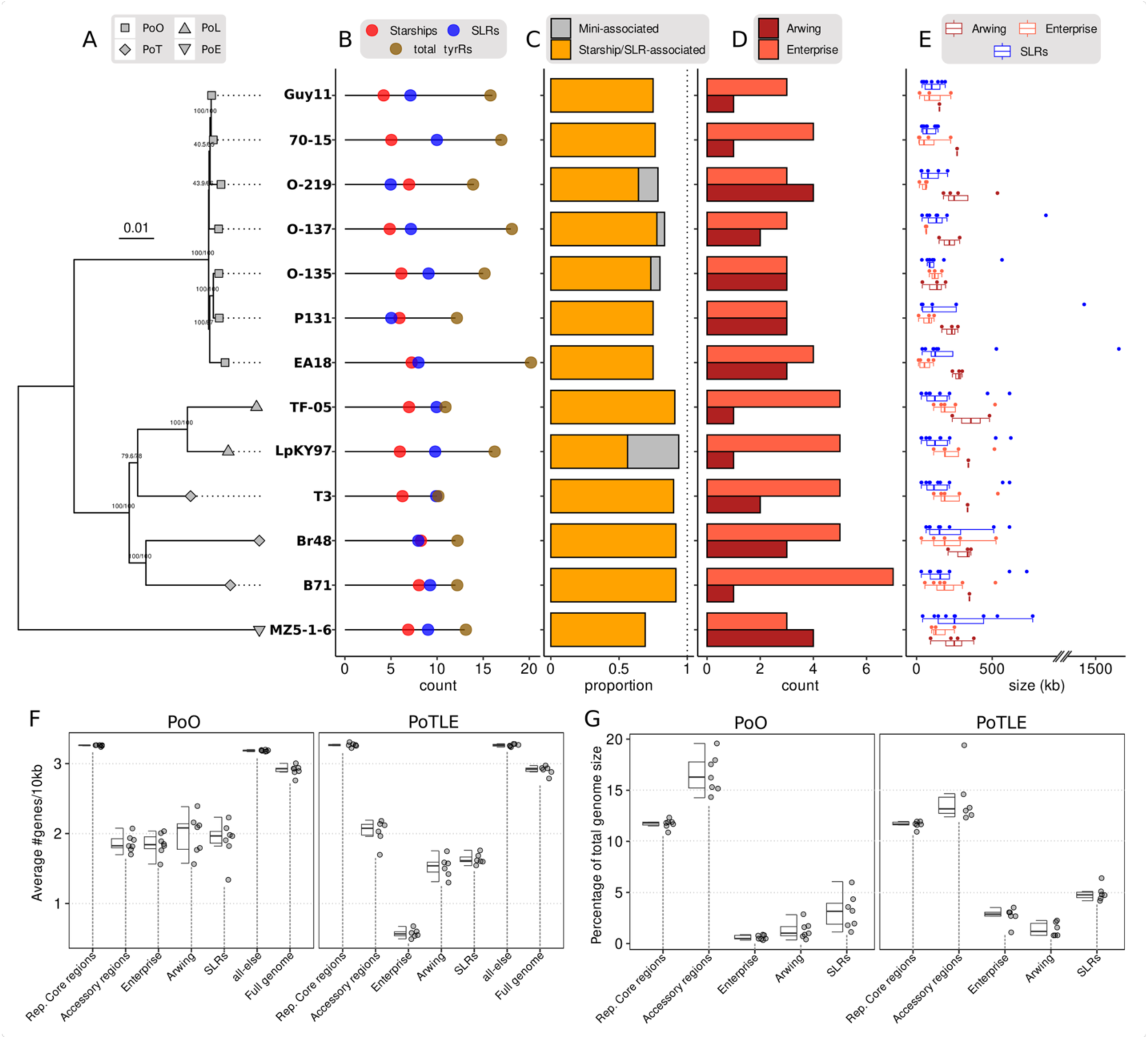
Diverse *Starships* and *Starship-*like regions (SLRs) characterize the *Starship* compartment of *P. oryzae*. (A) Mid-rooted BUSCO-based maximum likelihood phylogeny of *P. oryzae* strains used in this study. Terminal node shapes denote pathotype assignments. (B) Total count of Tyrosine Recombinases (*tyrRs*), SLRs and *Starships* detected in each assembly. (C) The proportion of total *tyrRs* associated with *Starships*/SLRs or mini-chromosomes. (D) The number of *Starships* with *tyrR* captains from either the Arwing or Enterprise families. (E) Sequence length variation of *Starships* from the Arwing and Enterprise family elements and SLRs. (F) The average gene density per 10kb and (G) percentage of total genome size for each defined compartment; Representative Core regions (a subset of +/- 50kb regions surrounding 50 randomly selected BUSCO genes), Accessory regions (all regions defined as large PAVs but not associated with tyrRs and therefore not considered to be *Starships* or SLRs), Enterprise family *Starships*, Arwing family *Starships*, SLRs, all-else (all other regions not contained within Core, Accessory, SLRs or *Starships*) and Full genome (the entire genome for each strain). Strains were split into the PoO and PoTLE lineages.

By integrating the outputs of starfish and stargraph, we provide the first compendium of *Starships* and SLRs, referred to as the *Starship* compartment, for *P. oryzae*. Additional *Starship-*derived regions may exist within our dataset (e.g. those that have lost *tyrR* sequences but have retained other *Starship* cargo), which are not considered here as SLRs, although this may be warranted in future analysis. To test our overarching hypothesis that the *Starship* compartment makes distinct contributions to fungal genome evolution, we defined two additional compartments for comparison. First, we defined a representative sample of the core compartment as a set of 50 randomly selected single copy BUSCO genes +/-50kb flanking sequence that are present in all 13 analyzed strains. We then defined the accessory compartment as the set of all large PAV regions (>=30kb) present in the pangenome variation graph that did not contain any predicted *tyrRs* and thus vary in their presence among the 13 strains.

Between four to eight *Starships* and between five to ten SLRs were identified in each genome (Fig. 1B, Fig. S2 and S3, Table S2-3). A phylogeny of the *Starship* captains confirmed that all elements belong either to the Arwing or Enterprise families, which represent two of the eleven previously defined phylogenetic families that capture the diversity of all known *Starship* elements (Gluck-Thaler and Vogan 2024; Vogan et al. 2021). There was a trend of more Enterprise family elements in the *Triticum* and *Lolium* pathotypes (PoTL) compared to the *Oryzae* pathotypes (PoO) (Fig. 1C, Table S3). Analysis focused on comparisons within and between *P. oryzae* pathotypes that infect rice (PoO) and those that infect *Triticum, Lolium,* and *Elusine* collectively referred to as PoTLE. These groupings were used for analysis because they reflect pathotype relatedness and variability (Fig. 1A) consistent with previous reports (Gladieux, Condon, et al. 2018; Peng et al. 2019; Rahnama et al. 2023). The identified *Starships* ranged from 12.8 kilobases (kb) to approximately 500kb, while SLR sizes were more variable and occasionally very large, with the two longest regions spanning 1400 and 1600kb (Fig. 1D). The overall gene density of both *Starships* and SLRs regions tended to be lower than representative core regions and similar to accessory regions (Fig. 1F, Table S4; see methods). The SLRs represented a larger percentage of the genome compared to *Starships*, and the median genome percentage of combined *Starship* and SLRs (i.e. the *Starship* compartment) was 4.3% for PoO strains and 9.2% for PoTL (*Triticum and Lolium*) strains (Fig. 1G, Table S5). This makes the *Starship* compartment an overall smaller but similar percentage of the genome than the accessory compartment, which accounted for a median of 16.6% of the genome for PoO strains and 14.2% for PoTLE strains (Fig. 1G, Table S4-5).

Mini-chromosomes in *P. oryzae* represent particularly interesting and relevant regions of the genome because they mediate horizontal genetic exchange and display significant sequence variation (Peng et al. 2019; Langner et al. 2021; Liu et al. 2024; Lin et al. 2025; Fang et al. 2025), raising the question of how they interact with *Starship* elements. However, our annotation methods could not confidently identify *Starship*s or SLRs on mini-chromosomes present in our analyzed strains. This is because mini-chromosomes typically have high sequence divergence, making it difficult to find clearly segregating *Starship* insertion sites. However, there are *tyrR* coding sequences present on mini-chromosomes, which may indicate that *Starships* have inserted into mini-chromosomes at some point in the past, but their association may also be due to other sequence exchange events. For most subsequent analyses, we therefore focused on core chromosomes with reliable *Starship* and SLR annotations and excluded mini-chromosomes.

### *Starships* in the Arwing family display hallmarks of mobility in *P. oryzae* genomes

Genomic mobility is a key feature of *Starships* and is mediated by the *tyrR* captain (Urquhart et al. 2023). To understand the impact of *Starship* mobility on *P. oryzae* strain variability, we further classified all *Starships* into navis groups (plural “naves”) using amino acid-based orthologous relationships between *tyrR* genes, as previously described (Gluck-Thaler and Vogan 2024). We then classified all *Starships* into haplotypes based on pairwise *k*-mer similarity clustering across their entire nucleotide sequences, resulting in a joint navis-haplotype designation for each *Starship* that was used to discriminate between different elements (methods). We assigned charismatic names to all element naves with evidence of mobility (i.e. where the *tyrR* captain from a given navis was present at >1 distinct genomic site across the 13 strains), resulting in the *Cloud*, *Arrow* and *Wolfen* naves. All three of these naves with evidence of mobility at the population level are members of the Arwing family. All other naves retained their randomly assigned navis number.

We then designated *Starships* as stable, variable, or unique based on their segregation frequencies at insertion sites and their navis-haplotype identifier (Table S2). Stable *Starships* have a single haplotype and are present at the same syntenic locus across all strains of a given pathotype. Variable *Starships* are present at syntenic loci in at least two (but not all) strains of a pathotype, while unique *Starship* insertions are present at a single genomic locus in only one strain. In the PoO strains, just over 50% of the *Starships* in the Arwing family are unique (7/13) and no single element in this family is stable (Fig. 2, Table S2). In contrast, Enterprise family elements are unique 9% of the time (2/23) and 60% stable (14/23). In the PoTL strains, 38% of the Arwing family elements are unique (3/8) and none of them are classified as stable. Enterprise family elements in PoTL strains are only 7% unique (2/27) and 78% stable (21/27). These observations indicate high stability for the Enterprise family elements across all strains, but contrasting stability for Arwing family elements across pathotypes (Fig. 2, Table S2). Differences between pathotypes could be due to the recent emergence of wheat blast (PoT) strains, and their shared lineage with PoL strains (Rahnama et al. 2023). There are many potentially unique elements in strain MZ5-1-6, which belongs to the PoE pathotype, but these were not categorized as stable or unique because this was the only PoE strain analyzed. These results show contrasting stability profiles between the two *Starship* families, which may indicate differences in activity, selection, or time since insertion.

**Figure 2.**
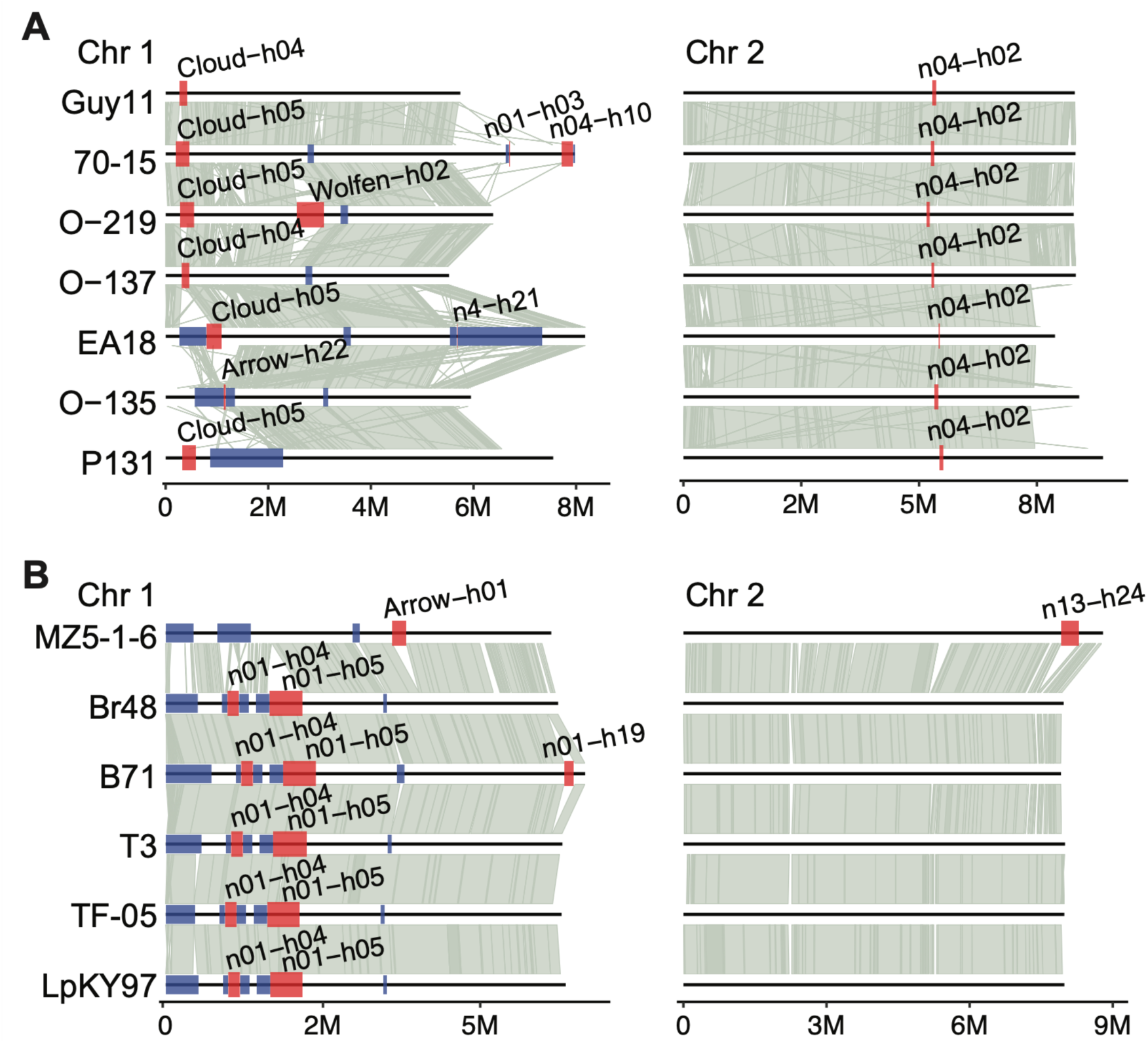
*Starships* in the Arwing family are highly variable across *P. oryzae* strains. (A) Chromosome synteny plot for PoO pathotype strains labeled to left (chromosome 1 left, chromosome 2 right; see supplemental figures 2 and 3 for other chromosomes). Chromosome length shown at bottom in millions of base pairs. Red and blue boxes show the location of *Starships* and SLRs, respectively. *Starships* are labeled above each element as navis-haplotype. Grey links between chromosomes indicate syntenic regions defined as greater than 90% similarity greater than 5kb. (B) Chromosome synteny plot for PoTLE strains depicting *Starships* and SLRs on chromosomes 1 and 2 as shown in A.

### *Starships* in clonal PoO strains are hotspots of genomic variation

We next performed a network analysis to begin quantifying and comparing the levels of sequence diversity in the *Starship* compartment to the core and accessory compartments to test our hypothesis that *Starships* make distinct contributions to overall genomic variation (Fig. 3, Fig. S4; Table S6-8). Each node in the networks corresponds to a particular region, and edges between nodes represent their pairwise nucleotide *k*-mer based Jaccard similarity across their entire sequence. Clusters within the network were defined as groups of n≥2 nodes connected by one or more edges. Network clustering largely recapitulated *Starship* navis classification based on *tyrR* orthology, meaning that *Starships* assigned to the same navis generally had high cargo sequence similarity which we would expect if these elements shared a recent common ancestor as indicated by the similarity of their *tyrR* captains (methods). For example, all *Starships* from the *Cloud* navis (Arwing family) were found in a single cluster with only one additional SLR that is likely derived from a *Cloud Starship* (Fig. 3C). Of the nine clusters identified in PoO strains, five (56%) were composed of more than one type of *Starship*-associated region, namely, some combination of an Arwing family element, an Enterprise family element, or SLR (Fig. 3C). For the 13 PoTLE network clusters, only two (15%) were composed of more than one type of *Starship-*associated region (Fig. 3H). Across all *P. oryzae* strains, mixed clusters most commonly contained Enterprise family elements and SLRs. The remaining elements were present in small, highly connected clusters of a single element navis-haplotype, or as singletons with no connections (Fig. 3C, H).

**Figure 3.**
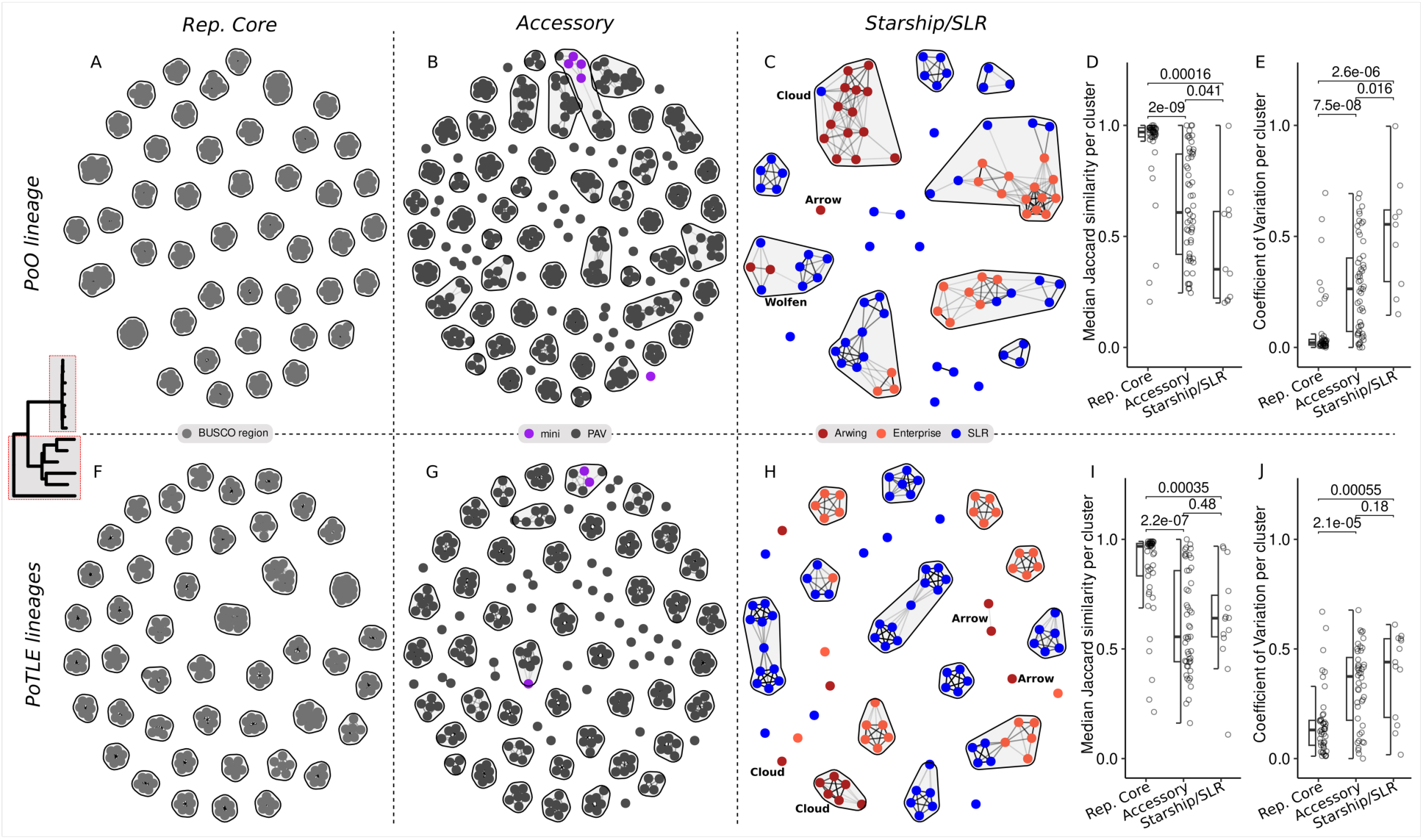
The *Starship* compartment of PoO strains contains the most variable regions of the genome. Comparisons of homologous DNA region (i.e. cluster) variability across genomic compartments for the PoO (A-E) and PoTLE lineages (F-J). Network nodes represent genomic regions, and edges are based on pairwise *k*-mer Jaccard similarity. Clusters of homologous regions were defined as any group of interconnected nodes. (A, F) Core genomic regions consist of +/-50kb regions surrounding 50 randomly selected BUSCO genes. (B, G) Accessory genomic regions are the set of present/absent variable (PAV) regions, but do not contain *Starship*-associated *tyrRs*, as defined within the PoO lineage (B) or the PoTLE lineages (G) based on the genome graph method. (C, H) The *Starship* compartment, composed of *Starships* and *Starship-*like regions (SLRs), from the PoO lineage (C) or the PoTLE lineages (H) where each node is colored according to its classification as either an Arwing family element (brown), Enterprise family element (salmon) or SLR (blue). Nodes and clusters representing specific Arwing family elements (*Arrow*, *Cloud*, *Wolfen*) are labelled. (D, I) Median Jaccard similarity per cluster for the three categories of genomic elements listed on the X-axis. (E, J) Coefficient of variation per cluster for the three categories of genomic elements listed on the x-axis. Individual regions are depicted as open circles and summarized as box and whisker plots depicting the interquartile range and median shown as a thick black line. Wilcoxon test p-values are shown above individual pairwise comparisons.

Median *k*-mer based Jaccard similarities were calculated per cluster to further understand how sequence diversity is influenced by genomic compartment. The within-cluster Jaccard similarity was highest for the core compartment, and significantly lower within the PoO *Starship* compartment (p=0.00016; Fig. 3D). Likewise, the PoO *Starship* compartment had a significantly higher coefficient of variation compared to the other two compartments (p<0.016; Fig. 3E), indicating that DNA segments in the PoO *Starship* compartment differ more from each other compared to segments within the other two compartments. The significant differences in both the median Jaccard similarity and coefficient of variation remained when comparing the accessory compartment with the *Starships* alone but not SLRs alone (Fig. S5). In contrast, the PoTLE strains did not have a significant difference between accessory and *Starship* compartments, but both compartments were significantly different than the core compartment (Fig. 3I-J).

*Starship* analyses in other fungi have largely focused on transposition, transfer, and characterization of specific cargo phenotypes, but less is known about how *Starships* influence the overall trajectory of genome diversity. To understand if the elevated sequence variation identified in PoO *Starships* is common, we performed a similar *Starship* and SLR annotation and network analysis using 13 *Aspergillus fumigatus,* another model fungal species from a different taxonomic class whose *Starship* compartment we recently described (Gluck-Thaler et al. 2025) (Fig. S6). On average, we identified a *Starship* compartment made up on average of 770kb, the equivalent of 2.7% of the total genome. We identified a total of 15 clusters of *Starships* and SLRs, similar to the total found across the 13 *P. oryzae* strains (Fig. S6; Table S9). There was a greater overall richness of *Starship* families in *A. fumigatus*, with a total of six families compared to the two in *P. oryzae.* To understand how family diversity is related to the overall sequence diversity present in the genomes analyzed, we calculated the pairwise average nucleotide identity (ANI) between assemblies within each species separated by pathotype groups (Table S10). The lowest ANI, indicating the greatest sequence divergence, was found for the PoTLE strains, which is likely due to the mix of host infecting pathotypes, while the PoO strains had the highest median ANI, consistent with previous descriptions of very low diversity across the global clonal lineages of rice infecting *P. oryzae* (Fig. S6D) (Gladieux, Ravel, et al. 2018; Zhong et al. 2018). Notably, the ANI of the PoO strains was statistically higher than that of *A. fumigatus* (median of 99.79% vs 99.67% respectively) (Fig. S6D). The median Jaccard similarity of *Starships* in PoO and *A. fumigatus* was 0.37 versus 0.77 (p=0.038; Fig. S6E), and of SLRs was 0.54 versus 0.72 (p=0.18; Fig. S6E). The median coefficient of variation of *Starships* in PoO and *A. fumigatus* was 0.56 versus 0.25 (p=0.113; Fig. S6F), while in SLRs was 0.5 versus 0.5 (p=0.81; Fig. S6F). These results show that *Starships* in PoO strains are significantly more sequence diverse than in *A. fumigatus*, despite the *A. fumigatus* strains having higher background genome-wide sequence diversity. These results point to the possibility that *Starships* differ in their contributions to genomic diversity depending on the fungal species in question and may be a particularly substantial source of genomic diversity in lineages such as PoO that reproduce clonally.

### Large structural variants characterize the mode of sequence diversification in *Cloud Starships*

We next sought to identify an origin and potential mechanism responsible for the elevated levels of genetic variation observed in some *Starship* sequences. To follow up on our observation that the *Starship* compartment is significantly more diverse at a sequence level compared with the core and in some cases the accessory compartment (Fig. 3), we compared the relative frequencies of all SNPs and insertions/deletions (INDELs) from pairwise alignments of *Starship* sequences within the same network clusters (Fig. 4C-D, Table S11). Remarkably, we found that *Cloud Starships* (cluster size n=21) have significantly more sequence impacted by medium (1-10kb) and large (>10kb) INDELS compared with SNPs. To understand if these results were specific to the *Cloud* elements or generally true for all other *Starships* and all other compartments, we applied the same calculations to the second largest network cluster containing Enterprise family elements (n= 18), the two largest comparable clusters from the accessory regions network (n= 26 and 19) and the representative core region clusters (n= 650) (Fig. 4D, S5, Table S11). We only detected a significantly greater amount of sequence impacted by INDELs <1kb in the Enterprise family cluster and representative core regions (Fig. 4D, Table S11). Otherwise, no significant differences were detected between the total amount of sequence impacted by any INDELs versus SNPs in any other genomic compartment, indicating the results for the *Cloud* elements are unique We manually confirmed the prevalence of multiple independently segregating large INDELs in *Cloud* elements by creating synteny alignment plots of representatives for each *Cloud* haplotype (h1-h7) (Fig. 4B). This included *Cloud* h4 and h5 that are largely syntenic except for the large insertion in h5, which is likely due to the nested insertion of another *Starship* considering the presence of a *tyrR* being the closest gene to one edge of the insertion. Aside from this example, the other changes to cargo content show no evidence of being caused by nested insertion. Additionally, these regions appear to contain novel cargo such as two uncharacterized biosynthetic gene clusters (BGCs) in h2 and h6 (Fig. 6B; Fig. S7). In contrast to the gain and loss of unique sequences contributing to large INDEL formation, one region stood out as the only example of genes being duplicated within different haplotypes (Fig. 6B; Fig. S7). This duplicated region contained two genes annotated as N-terminal amidases (ENOG503P0F8), proteins previously described as virulence factors in diverse pathogens (Gardiner et al. 2012; Yang et al. 2018). The same genes are not annotated in *Cloud*-h1 as both are heavily truncated due to nonsense mutations resulting in premature stop codons. As the region containing these genes is absent from other *Cloud* haplotypes, this suggests that the gain, loss, and duplication of these genes has occurred all within *Cloud* haplotypes. These element-wise comparisons demonstrate an extreme case of sequence diversity within the *Cloud* elements from the Arwing family, characterized by large INDELs that result in the gain, loss, and duplication of genes with predicted roles in biotic interactions.

**Figure 4.**
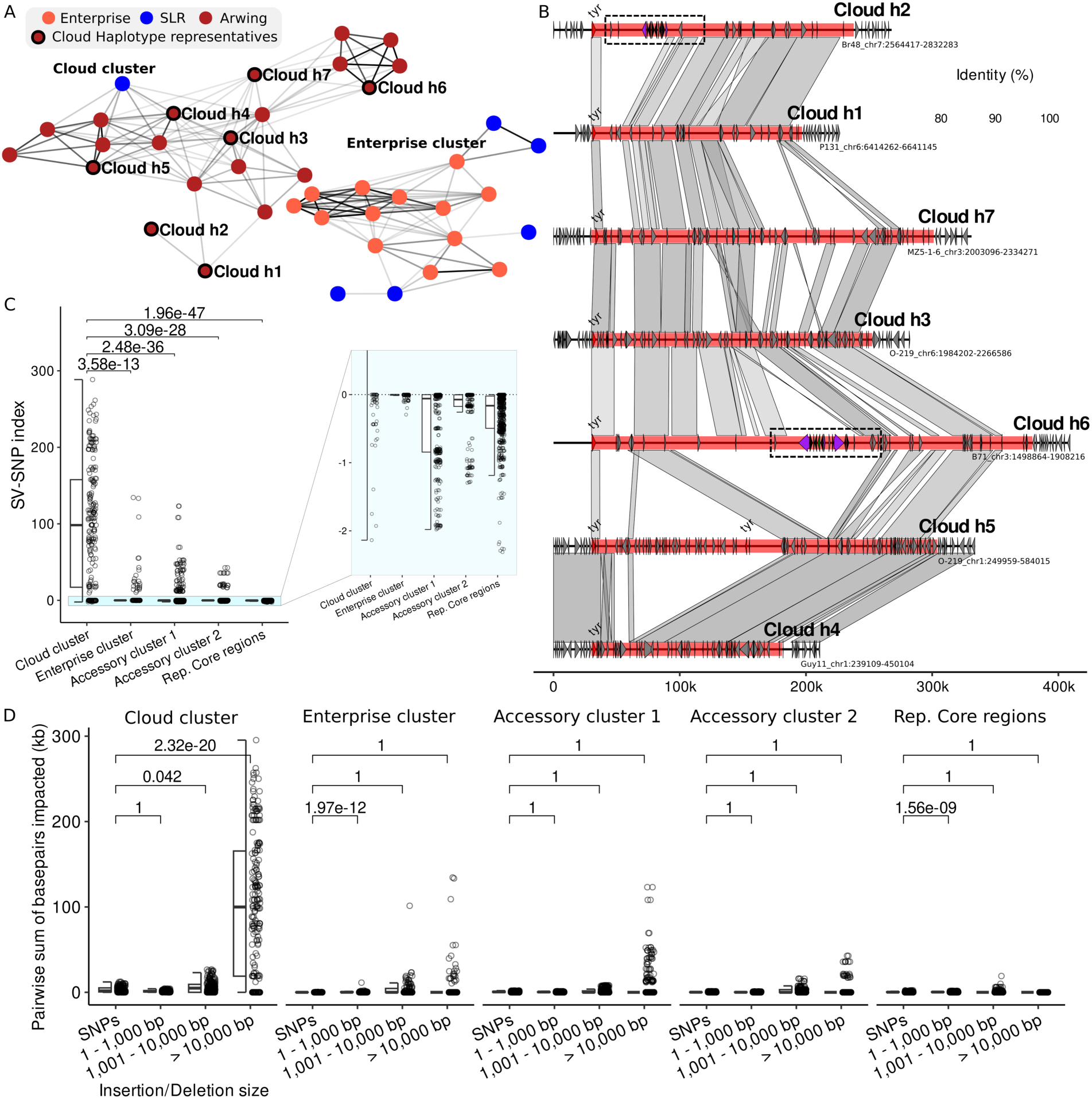
*Cloud Starships* contain multiple independent insertion, deletion, and duplication structural variants across strains and are hotspots for genomic variation. (A) Subsets of the two largest clusters from the *k*-mer-based Jaccard similarity network. Clusters were defined as any group of interconnected nodes and represent homologous genomic regions. (B) Synteny plots of representative elements for each *Cloud* haplotype with 30kb flanking regions around each element. Putative BGCs are highlighted by dotted boxes in *Cloud*-h2 and -h6. (C) The summed total number of nucleotide bases impacted by Insertions/Deletions (INDELs) greater than 10kb minus the number of SNPs (i.e. the SV-SNP index (kb)) from different compartment clusters. The pale blue highlighted region and corresponding enlarged region helps visualize the complete distribution of the SV-SNP index values. Positive SV-SNP values indicate a greater proportion of bases impacted by large INDELs, while negative SV-SNP values indicate a greater proportion impacted by SNPs. P-values were calculated using one-sided Wilcoxon tests with FDR correction for multiple testing to evaluate whether the SV–SNP index is significantly greater in the *Cloud* cluster. (D) The summed total number of base pairs impacted by mutations in pairwise comparisons of all clusters from different genomic compartments (*Cloud* cluster, Enterprise family cluster, the two largest clusters from the **a**ccessory regions and **r**epresentative **c**ore regions). INDELs were binned into sizes of 1-1000bp, 1001-10000bp, and greater than 10kb. P-values were calculated using one-sided Wilcoxon tests with FDR correction for multiple testing to evaluate whether the sum of bases impacted by SNPs was less than each INDEL category.

To further compare the impacts of large structural variants across the various compartments, we compared the total number of base pairs impacted by INDELs >10kb minus the total number of base pairs impacted by SNPs (which we define as the SV-SNP index) within each cluster. Higher positive values of this index indicate a greater proportion of sequence impacted by large INDEL mutations, whereas smaller negative values indicate a greater proportion of sequence impacted by SNP mutations. Both the *Cloud* cluster and Enterprise family cluster had significantly greater SV-SNP indices (median: 94714bp (p= 1.96E^47) and –3bp (p=1.22E^30) respectively) compared to the core compartment clusters (median: −163bp; Fig. 4C), indicating that large structural variants drive sequence evolution within *Cloud Starships* and to some extent the *Starships* compartment as a whole. Both the Cloud and Enterprise family clusters also have significantly larger SV-SNP indices compared to both Accessory compartment clusters (median: −59bp (p<3.09E^28) and –73bp (p<6.33E^12) for Accessory clusters 1 and 2, respectively). Although a single large INDEL by definition impacts a larger number of base pairs than a single SNP, we observe significant variation in the SV-SNP index across the various compartments, indicating that the enrichment of large INDELs observed within *Cloud Starships* is neither a statistical nor tautological artifact. We further manually confirmed that the total number of base pairs impacted by large INDELs arises from many independent mutations and not simply a single large mutation (Fig. 4B). Together, these genetic variant analyses indicate that sequence evolution within *Cloud Starships* is driven to a greater extent by mechanisms generating large structural variants that result in extensive gene gain, loss and duplication, whereas sequence evolution in the core and accessory compartments is driven primarily by single base pair mutations.

### *Starships* are associated with elevated rates of presence/absence variation in virulence associated genes involved in plant-pathogen interactions

Given our observation that *Starships* contribute to overall sequence diversity in *P. oryzae*, we next checked if known effectors and virulence coding sequences were associated with the *Starship* compartment. To this end, the locations of 116 virulence-associated coding sequences was determined in the 13 *P. oryzae* genomes, including representatives of 94 *Magnaporthe* Avrs and ToxB-like (MAX) effector ortholog groups (Le Naour—Vernet et al. 2023) and 22 experimentally characterized avirulence (Avr) and infection-related sequences (Table S12-13). These coding sequences were assigned to the *Starship* compartment if they were located on a *Starship* or SLR. Although the majority of universally conserved MAX and Avr coding sequences with no PAV were not associated with the *Starship* compartment, we identified a total of 16/94 MAX and 7/22 Avrs and virulence associated coding sequences in the *Starship* compartment in one or more strains (Fig. 5 A-B).

**Figure 5.**
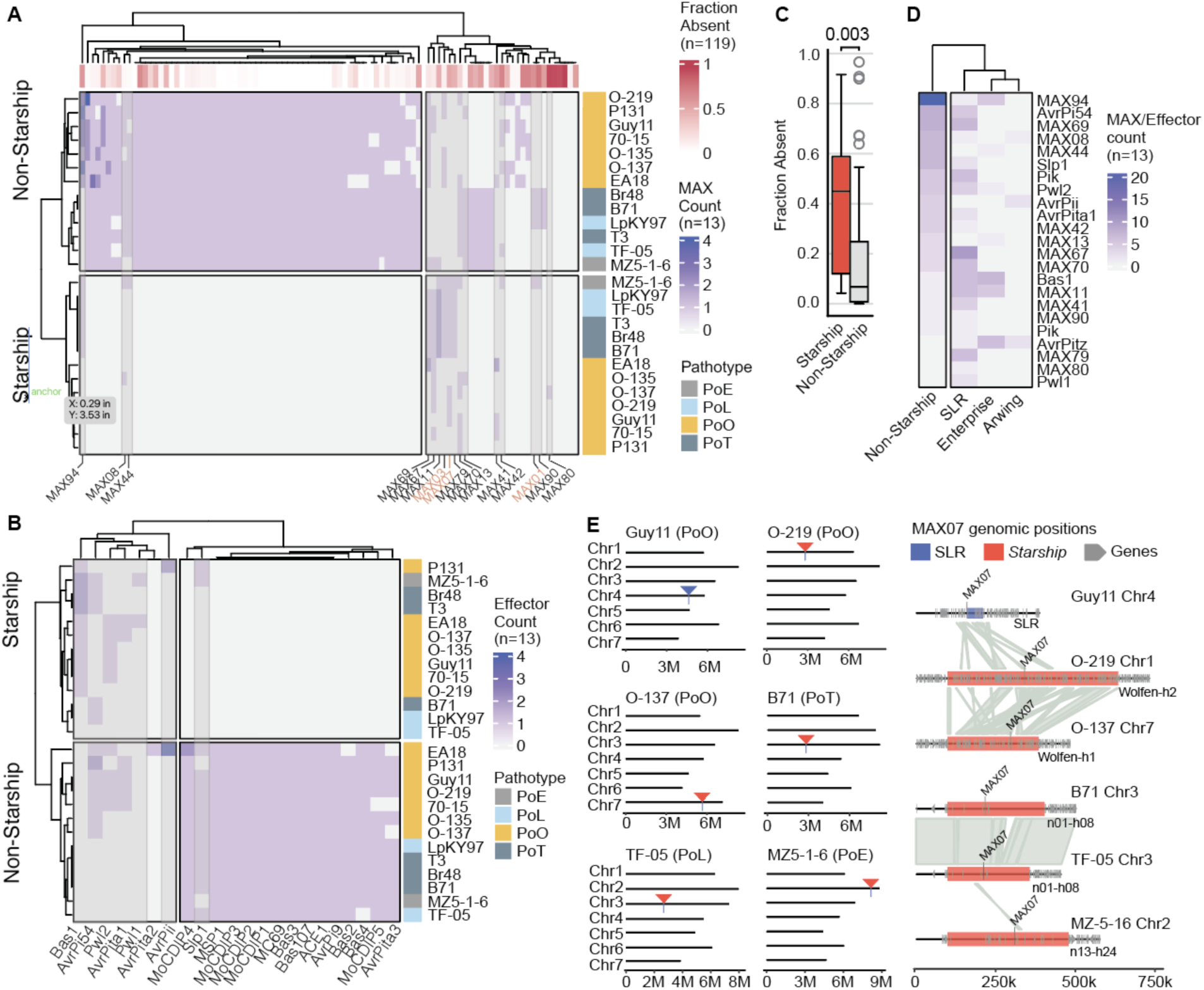
Effector coding sequences in the *Starship* compartment have high interstrain variability and mobility. (A) Heatmap of MAX effector coding sequences in 13 *P. oryzae* genomes colored by pathotype. K-means clustering (k=2) was applied to MAX effectors (columns) and strain regions (rows) that either overlapped the *Starship* compartment or were labeled non-*Starship*. Each strain has two rows: one summarizing across all non-*Starship* regions and another summarizing across all *Starship* elements. MAX coding sequences that overlap at least one *Starship* or SLR in one strain are labeled along the bottom and highlighted with a grey box. The fraction of strains from a larger population (n=119) missing a MAX coding sequence is reported as Fraction Absent along the top of the heatmap, as previously reported (Le Naour—Vernet et al. 2023). MAX sequences in the *Starship* compartment that have an Avirulence name are highlighted in orange and not shown in B. (B) Heatmap of characterized Effector and Avirulence coding sequences in the *P. oryzae* strains. The heatmap is organized as described in A and excludes any Avr sequences that overlap with MAX groupings. (C) Boxplot of the Fraction Absent for MAX effector coding sequences summarized in *Starship* (red) versus non-*Starship* (grey) genomic regions from the set of 119 strains. Statistical significance tested using a two-sided t-test, p-value = 0.003. (D) Heatmap of MAX effector and avirulence coding sequences located in the non-*Starship* regions, or in an SLR or *Starship* across the 13 *P. oryzae* genomes, broken up by *Starship* family. (E) Chromosome locations of MAX07 (*AvrPiz-t*) coding sequence in six representative genomes across all pathotypes (right). A synteny plot for the individual loci from the six genomes (left). The MAX07 coding sequence is labeled, along with other annotated coding sequences depicted as gray arrows. The coordinates of SLRs and *Starships* are depicted by color boxes. Grey links indicate syntenic regions defined as having >90% nucleotide similarity across >2 kb.

To determine the patterns of *Starship-*associated effector PAVs at a larger population level, we examined a previously published MAX effector PAV analysis calculated across a diverse collection of 119 *P. oryzae* strains (Le Naour—Vernet et al. 2023). The 16 MAX ortholog groups that we identified in the *Starship* compartment had a statistically significantly higher PAV rate across the 119 strains compared to the rest of the MAX ortholog groups (median 45% versus 7% absent, two-sided t-test, p=0.003) (Fig. 5C). The association between MAX effectors in the *Starship* compartment and PAVs is likely underreported because five MAX ortholog groups with PAVs were not identified in any of our 13 long-read assemblies and had been classified as non-*Starship*. Three of these families (MAX68, 92, 93) have greater than 90% absence in the 119 strain dataset (Le Naour—Vernet et al. 2023), similar to MAX80 that had 92% absence and was identified on a *Starship* in PoO strain O-137 but not found in any other assembly. We anticipate that coding sequences belonging to at least these three highly variable MAX ortholog groups will be identified on *Starships* in future assemblies of new strains. The majority of MAX and Avr coding sequences were present on SLRs, followed by Enterprise family elements (the least active *Starship* family), and the least were present on Arwing family elements (the most active *Starship* family; Fig. 5D).

We also identified *Starships* as a potential mechanism contributing to variation in effector chromosomal location (i.e. genome position variation). For example, the MAX07 coding sequence was identified in 9/13 strains, but was present at five unique genomic locations across these strains (Fig. 5E). Specifically, the sequence was present on a stable Enterprise family *Starship* in all five PoTL strains, on a unique unnamed Arwing family element in MZ5-1-6 (PoE), on two separate chromosomes on two haplotypes of the *Wolfen Starship* (Arwing family) in O-219 and 0-137 (PoO), and on an SLR in Guy11 (PoO) (Fig. 5E). Importantly, MAX07 is part of the same orthogroup as MAX06 (Le Naour—Vernet et al. 2023) and the two comprise one structural group characterized by the effector *AvrPiz-t* (Lahfa et al. 2024). The AvrPiz-t protein targets rice E3 ubiquitin ligases to subvert pattern recognition receptor immunity (Li et al. 2009; Park et al. 2012; 2016), demonstrating critical virulence proteins as mobile *Starship* cargo. Thus, *Starships* not only impact effector PAV at a population level, but that they also mediate genome position variation which we speculate could result in variation in how these effectors are inherited and expressed.

To further examine the impact of *Starships* on effector-mediated *P. oryzae* virulence, we analyzed the presence and location of Nudix (nucleoside-diphosphate linked to moiety X) coding effectors that promote virulence in diverse pathogens (Dong and Wang 2016). In *P. oryzae*, secreted MoNUDIX hydrolyzes plant inositol pyrophosphate, triggering plant phosphate starvation signaling that subsequently reduces plant defense (McCombe et al. 2025). Three Nudix coding sequences were reported in PoO strain 70-15, two *MoNUDIX-1* sequences and one *MoNUDIX-2* (McCombe et al. 2025). We therefore checked the presence and location of Nudix coding sequences in our set of 13 strains. All seven PoO strains have one copy of a *MoNUDIX-1* sequence at a syntenic region toward the 3’ end of chromosome 6 (Fig. S8A). Four PoO strains have a second copy of *MoNUDIX-1* that is adjacent to an Enterprise-family *Starship,* which is in an annotated SLR in three of these four strains (Fig. S8B). The other three PoO strains are missing both the second copy of *MoNUDIX-1* and the *Starship-*SLR associated region, suggesting the second copy is associated with the presence of this *Starship* (Fig. S8B). Support this, the PoE and PoL strains contain one *MoNUDIX-1* coding sequence at a similar region on chromosome 6, but none of the PoTLE strains contain the second copy of *MoNUDIX-1* nor the *Starship-*SLR associated region (Fig. S8C). All 13 *P. oryzae* strains contain the *MoNUDIX-2* coding sequence. These results associate *Starship* variation with the increased copy number of *MoNUDIX-1* in PoO strains. Collectively, our data on *Starship-*effector associations suggest that *Starships* have impacted the evolution of well-characterized virulence coding sequences in *P. oryzae* by mediating copy number variation, PAV, and chromosomal location variation, which we speculate has a direct impact on the evolution of host-pathogen interactions and strain success in the field.

### Unique *Starship* insertions in *P. oryzae* were horizontally transferred from closely related species

Increasing experimental and population genomic evidence suggests *Starships* mediate horizontal gene transfer (HGT) both in the lab and in natural fungal populations (O’Donnell et al. 2025a; Bucknell et al. 2024; Urquhart et al. 2025). We therefore examined all *P. oryzae* elements for signatures of HGT against a large database of 10,010 publicly available fungal genome assemblies (methods). For the initial HGT screen against this large database, we extracted a set of candidate target genomes based on the presence of large alignable regions to our set of query *Starships* that exceeded expected thresholds for genome-wide sequence similarity. We then followed up the initial screen with a more refined “BLAST all” analysis, where all predicted nucleotide gene sequences from the query *P. oryzae* genomes were BLAST searched against a candidate target genome to obtain a range of sequence similarity values for all genes within the *Starship* in addition to an expected genome-wide background threshold (Urquhart et al. 2022). We found evidence for multiple horizontal transfers of *Cloud Starships* between *P. oryzae* and four other species within the *Pyricularia* genus (Fig. 6A-D; Fig. S9-S14 Table S14). While it is challenging to determine the exact donor and recipient lineages of these transfers, the unexpectedly high sequence similarity of these *Starships* across the different species (median 98.6% (98.1-99.7) pairwise similarity *Starships* vs median 90.7% (86.9-92.1) pairwise similarity of the Representative Core region) suggests the transfers occurred relatively recently. As expected, signatures of HGT within the *Starship* compartment stand out, as genes from the *Cloud Starships* have significantly greater signatures of HGT compared with the overall distribution of genes from the accessory and representative core regions (Fig. 6B).

**Figure 6.**
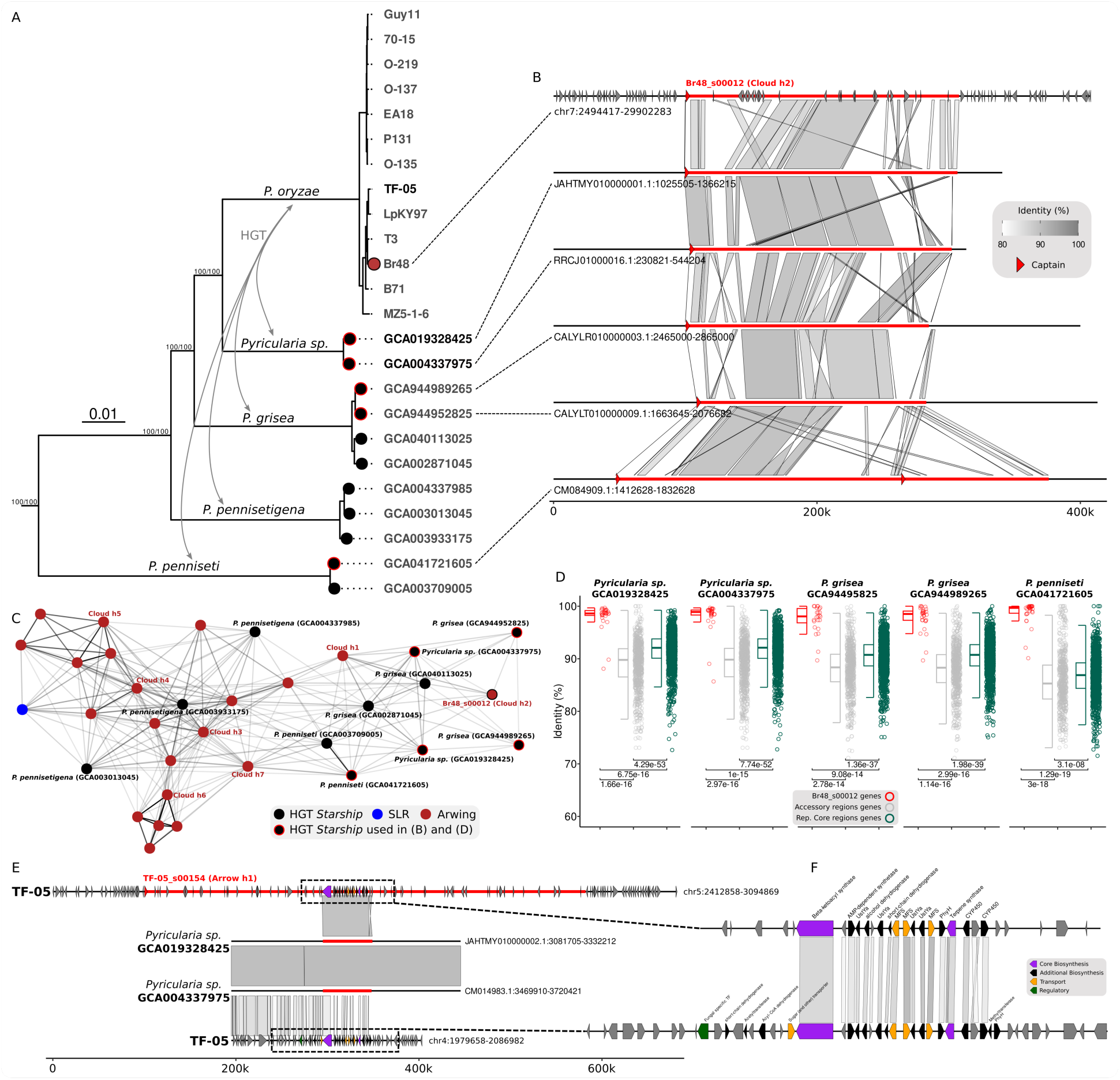
*Cloud Starships* have undergone frequent horizontal transfer across the *Pyricularia* genus. (A) A maximum likelihood (ML) tree made using 3291 BUSCO proteins of 13 *P. oryzae* assemblies and 11 assemblies from other species with evidence of a horizontally acquired *Starship*. The tree was rooted using *Gaeumannomyces* as an outgroup. Branch support is shown using 1000 Ultrafast Bootstraps and the SH-like approximate likelihood ratio test. Bold strain names/assembly accessions indicate those used in example (E). (B) Synteny alignments of select regions with signatures of HGT from representative assemblies across the genus. (C) The *Cloud Starship* cluster from a *k*-mer based Jaccard similarity network of all *Starships*/SLRs and regions with signatures of HGT from all genomes included in the phylogenetic tree. (D) Distribution of nucleotide identity values between all predicted gene sequences from strain Br48 and nucleotide sequences from candidate HGT donor/recipient genomes in bold. Br48 genes were grouped by whether they were located within the accessory compartment, core compartment, or within the *Cloud* haplotype 2 representative (Br48_s00012). Wilcoxon tests were used to compare the difference in identity between genes within Br48_s00012 and the **c**ore compartment with FDR corrections for multiple tests. (E) Alignments showing an example of a *Starship*-mediated biosynthetic gene cluster (BGC) transfer from *Pyricularia* sp. to *P. oryzae* TF-05 of a core BGC already present in *P. oryzae* TF-05 displayed in the bottom contig. (F) Protein sequence alignments between the *Starship-*transferred and core version of the BGC in TF-05.

To determine how sequence variability among the horizontally transferred *Cloud Starships* compares to our previous assessments, we repeated our network and synteny analyses to now include the horizontally transferred *Cloud Starships* from other *Pyricularia* species. We find that horizontally transferred *Starships* show similarly elevated levels of sequence variation as previously observed within the *P. oryzae*-specific *Cloud* cluster (shown previously, Fig. 4B-D). These results suggest *Cloud Starships* are not just hotspots of genomic variation in *P. oryzae* but potentially also in other *Pyricularia* species. Outside of the horizontally transferred *Cloud Starships*, we also observed the transfer of a large region of ∼50kb contained within an *Arrow* element to an unnamed, but closely related *Pyricularia sp.* (NCBI assembly accessions GCA019328425 and GCA004337975) (Fig. 6E). This region contains a near-duplicate copy of a BGC (Fig. 6F) whose paralog appears to be conserved within both *P. oryzae* and *Pyricularia sp..* Notably, the BGC paralog in the *Starship* is highly similar to that of the *Pyricularia sp.* copy and not the *P. oryzae* core chromosome copy, suggesting the additional BGC copy in *P. oryzae* is not a paralog, but was acquired by a *Starship* in *Pyricularia sp.* and then horizontally transferred (Fig. 6E-F).

### Arwing family elements are more euchromatic and their horizontal transfer does not always result in epigenetic silencing

The epigenome provides an important layer of genomic defense to suppress mobile DNA activity (Allshire and Madhani 2018). *Starship* analysis in *V. dahliae* identified similarly high levels of H3K27me3 in *Starships* and other accessory regions (i.e. adaptive genomic regions) (Sato et al. 2025). However, that analysis did not look at the presence of H3K9me3 to help distinguish types of heterochromatin, and *Starship* family or element level analysis was not reported to distinguish variation across elements. To address this knowledge gap, we analyzed the presence of three histone modifications across all genome compartments: trimethylation at H3-Lys9 (H3K9me3) and H3-Lys27 (H3K27me3), and acetylation at H3-Lys27 (H3K27ac). Both H3K9me3 and H3K27me3 contribute to the formation of heterochromatin, a repressed chromatin state, while H3K27ac is associated with transcriptionally active euchromatin (Freitag 2017). We previously collected data for all three histone modifications for four PoO strains (Guy11, O-219, O-137, O-135) (Zhang et al. 2021; Rowe et al. 2023), and data for both heterochromatic marks (H3K9me3 and H3K27me3) were publicly available for PoT strain (Br48) (Kobayashi et al. 2023). Peaks for H3K9me3, H3K27me3, and H3K27ac were intersected with the coordinates of all *Starships,* SLRs, representative core and accessory regions of these genomes. The core regions had statistically higher levels of H3K27ac compared to the accessory regions, Enterprise family elements, and SLRs, but were not significantly different compared to the Arwing family elements (Fig. 7A). On the other hand, Arwing family elements and the core regions were significantly less covered by H3K9me3 peaks compared to the Enterprise family elements, SLRs, and accessory regions (Fig. 7A). The core regions had significantly less H3K27me3 compared to all other regions (Fig. 7A). However, Arwing family elements displayed a trend for reduced H3K27me3, evidenced by the lower median H3K27me3 coverage (11.8%) compared to Enterprise family elements (82.3%) or SLRs (69.3%) (Fig. 7A). These collective results show that Arwing family elements have a euchromatic epigenome profile most similar to the core genome, and Enterprise family elements and SLRs have a heterochromatic epigenome profile most like the accessory genome.

**Figure 7.**
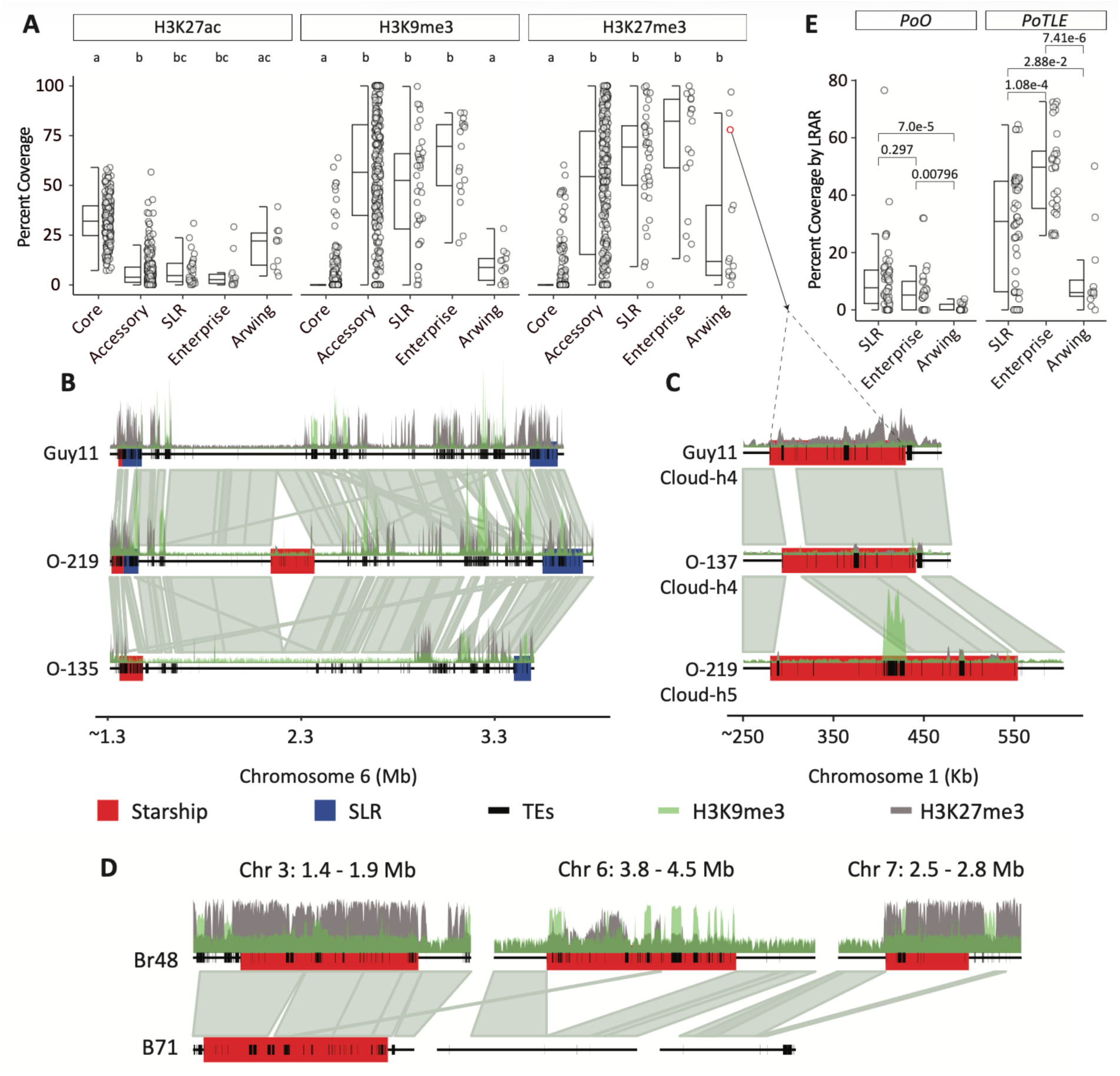
The Arwing family epigenome resembles that of the core compartment. (A) The percent coverage of representative core, accessory, SLR, and *Starship* regions for H3K27ac, H3K9me3, and H3K27me3 peaks. The H3K27ac plot does not include Br48. (B) Comparison of heterochromatic peaks in conserved Enterprise family elements, SLRs, and a unique Arwing element. Starship regions are highlighted by red boxes, SLR regions by blue boxes, and annotated transposable elements (TEs) by thick black lines. Locations of H3K9me3 and H3K27me3 histone modifications are shown as green and grey peaks, respectively. (C) Comparison of heterochromatic peaks in a *Cloud Starship* with variable epigenetic profiles across strains. Annotated elements are the same as described for B. (D) H3K27me3 and H3K9me3 associated with the three Arwing family *Starships* in the Br48 genome compared to the syntenic region of the related strain B71. Annotated elements are the same as described for B. (E) Percent coverage of *Starships* and SLRs by Large RIP-Affected Regions (LRARs) by *Starship* family and pathotype.

To further understand epigenetic variability within the *Starship* compartment, we visualized a ∼2 Mb region that highlights the epigenetic differences between Arwing and Enterprise elements on chromosome 6 of Guy11, O-219 and O-135 (Fig. 7B). The two ends of the region contain SLRs and an Enterprise family element in all three strains and have high levels of H3K9me3 and H3K27me3 (Fig. 7B). Interestingly, in the relatively syntenic middle portion of the region, O-219 has an Arwing family element insertion devoid of the heterochromatic histone modifications, which may indicate a recent insertion of this element and a potential escape of the element from heterochromatization (Fig. 7B). Contrary to the overall epigenetic profile of Arwing family elements, three Arwing elements had considerably higher H3K27me3 levels (Fig. 7A, Fig. S15A). One of these H3K27me3 marked Arwing family elements is a variable *Cloud Starship* present on chromosome 1 in most PoO strains analyzed (Fig. 7A, Fig. S15A). The *Cloud*-h4 *Starship* present in Guy11 and O-137 are syntenic to the *Cloud*-h5 *Starship* in O-219, 70-15, EA18, and P131, apart from a ∼125 kb insertion in *Cloud*-h5 (Fig. 7C, Fig. S2). In Guy11, a majority of the *Cloud*-h4 *Starship* is marked by H3K27me3, while the highly similar elements in O-137 and O-219 are not (Fig. 7C). Sequence content of the elements cannot explain the difference in H3K27me3 levels, as the syntenic regions are >90% identical and have similar TE annotations. The two other Arwing family elements with elevated H3K27me3 were both *Cloud Starships* present in the MoT strain Br48 (Fig. 7A, Fig. S15A). One was a variable *Cloud*-h6 *Starship* on chromosome 3, and the other a unique *Cloud*-h2 *Starship* on chromosome 7 of Br48 (Fig. 7D).

A previous analysis of the Br48 genome reported five large DNA segments with high similarity to DNA from *P. pennisetigena* and *P. grisea*, regions that the authors concluded had undergone horizontal transfer with *P. oryzae* (Kobayashi et al. 2023). We find that the regions identified in Kobayashi et al. correspond to annotated *Starships* and SLRs in Br48 (Fig. S16). The authors concluded that horizontally transferred DNA was associated with repressive histone modifications in Br48, likely a response by the recipient genome to silence the mobile foreign DNA (Kobayashi et al. 2023). However, a third Arwing family element in Br48, a unique *Arrow*-h2 *Starship* on chromosome 6, had low H3K27me3 coverage similar to the other Arwing family elements analyzed in PoO strains (Fig. 7D-E, Fig. S15A). Therefore, while some horizontally transferred *Starships* in Br48 are associated with heterochromatic marks, it is not universally true for all Br48 Arwing family elements. Furthermore, the horizontally transferred Arwing family elements in PoO strains lack substantial H3K27me3. Collectively, horizontally transferred elements identified in *P. oryzae* are predominantly from the Arwing family and have a low association with H3K27me3, but variation in histone modification profiles between related elements is seen between strains. The mechanism and consequence of this epigenome variation is currently not known but does appear to be epigenetic, i.e. not DNA sequence dependent.

To understand what factors might be contributing to the varied epigenetic states within the *Starship* compartment, we compared transposable element (TE) content and signatures of repeat induced point mutation (RIP) across all elements. The TEs in all 13 strains were annotated using RepeatMasker and a comprehensive TE library generated from representative *Magnaporthe* genomes of each pathotype (Nakamoto et al. 2023). Arwing family elements had significantly lower TE content compared to Enterprise family elements (mean 10.4% vs 52.1% for Arwing and Enterprise respectively, one-sided t-test, p<2.2e-16) (Fig. S15B). However, the three H3K27me3 marked Arwing family elements did not have distinctly different TE content compared to the other Arwing family elements, suggesting that differences in histone methylation are not driven by differences in TE content. Comparisons of TE content between stable, variable, and unique *Starships* demonstrated that stable elements have the highest TE content, while variable and unique elements are statistically similarly lower (Fig. S15B). We used Large RIP Impacted Regions (LRARs) as defined by the RIP annotation software RIPper (Wyk et al. 2019) to measure the impact of RIP across the *Starship* compartment. Within the PoO lineage, a significantly lower number of total bases and percent genome were impacted by RIP compared to the PoTLE lineage (p=0.0012) (Fig. S17A-B) and a significantly greater amount of RIP present in the PoTLE lineage within each compartment except the Representative Core regions (p=0.014-0.00117) (Fig. S17D; Fig. 7E), consistent with previous reports (Zong et al. 2024). In the *Starship* compartment, the Arwing family elements are the least impacted by RIP (Fig. 7E; Fig. S17E). Compared to random genomic sampling, PoO strains show a large and significant enrichment of RIP only within the Accessory compartment (Fig. S17E). In contrast, PoTLE strains exhibit large and significant RIP enrichments not only in the Accessory compartment but also in SLRs and Enterprise elements (Fig. S17E). In both lineages, the core compartment had a significantly lower enrichment in RIP (Fig. S17E). These results show that RIP is overall not a substantial force impacting PoO strains, and regions affected by RIP reside in the accessory compartment. The PoTLE strains have relatively more signatures of RIP, which are also associated with the accessory compartment, along with Enterprise family elements and SLRs. The identified differences in epigenetic marker distribution and RIP signatures may play a mechanistic role in mediating the mobility and elevated diversity of the Arwing family, which includes the *Cloud* elements.

## DISCUSSION

The concept of genome compartmentalization integrates evolutionary and mechanistic understandings of the genome, but the accessory compartment requires further refinement because it currently includes many types of PAV regions with different mechanistic origins and maintenance mechanisms. Although *Starships* do create PAV regions, we argue that they warrant their own separate compartment due to their ability to transpose and horizontally transfer dozens of protein coding genes, unlike other PAVs. At a minimum, *Starship* activity thus provides a mechanism for the origin of specific types of PAVs including horizontally acquired genotypes and translocations via transposition. Here, we present a formal analysis of the *Starship* compartment in relation to the canonical core and accessory compartments (A. Urquhart et al. 2024). We consider all regions composed of *Starships* and SLRs as part of this newly defined compartment to comprehensively assess the impact of these giant transposons on genome evolution. By considering *Starships* as a distinct compartment, we are effectively refining the umbrella concept of the accessory compartment by partitioning those genomic regions whose evolution is directly shaped by *Starship* activity. A logical extension therefore is that other components of the accessory compartment, such as mini-chromosomes (e.g., dispensable and accessory chromosomes), should similarly occupy their own separate compartment carved out from the umbrella accessory compartment term because they too have distinct molecular and evolutionary properties. Additionally, horizontal transposon transfer of non-*Starship* elements was recent found to account for approximately 8% of fungal genome TE content on average, but this number varied significantly over fungal biodiversity (Romeijn et al. 2025). Distinguishing between the granular contributions of different compartments thus stands to improve our mechanistic understanding and descriptions of how genomic variation arises. As a first step, we tested the hypothesis that the *Starship* compartment makes distinct contributions to fungal genome evolution as compared to other genomic compartments in terms of variation driven by *Starship* abundance, transferability, mutability and epigenetic modifications.

Although only a few species have been systematically examined at a population level, a perhaps unsurprising emerging trend is that the total fraction of a given genome made up of the *Starship* component varies enormously among strains and species. A recent study of 56 strains from the *Verticillium* genus found that *Starships* consist of or have contributed to the formation of approximately half of known accessory genomic regions in reference isolates and consist of ∼2-5% of total genome sequence (Sato et al. 2025). Using genome graph-based methods, we previously found that upwards of 15% of a *Penicillium camemberti* var. *caseifulvum* strain’s genome was made up of *Starships* and SLRs (O’Donnell et al. 2025a). Applying those same genome graph methods to *A. fumigatus* here revealed that on average 2.7% of genome sequence among strains is made up of *Starships* and SLRs. In *P. oryzae*, we found that the *Starship* compartment on average comprises 7% of the genome, comparable to the average percentage of the genome taken up by mini-chromosomes (6.2%) in our analyzed strains. When taken together, this means that *Starships* and mini-chromosomes in *P. oryzae* on average account for approximately the same magnitude of genome sequence as the remaining canonical accessory compartment as defined by our genome graph PAV analysis. Our findings argue for similar comparative approaches across genomic compartments to be applied to other fungal species to better understand how the rates, prevalence, and relevance of these diverse mechanisms of genome variation vary or not across different lineages, and what their unique contributions may be to genome evolution.

A distinct feature of the *Starship* compartment in *P. oryzae* is its mutability. *Starships* are known hotspots of large genomic rearrangements in several fungal species: in some cases, they coincide with up to ∼1/3 of chromosomal translocations, inversions and fusions (Gluck-Thaler et al. 2022; Sato et al. 2025). Here, we show that specific types of *Starships* are themselves hotspots for another type of SV: large insertions and deletions >10kb that are not explained by smaller transposable element activity. Active elements in the *Cloud* haplotype appear to evolve primarily by means of large SVs, more so than that of any other *Starship* haplotype or region within the accessory compartment. Additionally, our HGT analysis identified *Cloud*-like elements in other *Pyricularia* species that are similarly impacted by SVs, suggesting this variation may not be entirely species specific but rather may be a property of the *Starship* itself, or of an interaction between the *Starship* and the genomic background of *Pyricularia* fungi. This is particularly notable as several population studies have found that large SVs are rare, primarily impact intergenic regions, and are in most cases due to the movement of small, individual transposable elements (O’Donnell et al. 2023; Schloissnig et al. 2025). In a large survey of SVs in the plant pathogen *Zymoseptoria tritici,* although they detected an enrichment in accessory regions, only 0.4% of INDELs were larger than 10kb (Badet et al. 2021). Similarly, in *Fusarium graminearum*, SVs greater than 1kb make up only 0.25% of the total INDELs identified genome-wide (Dhakal et al. 2024). The general rarity of large SVs in these prior studies resembles our findings of large SV depletion in *P. oryzae*’s *c*ore compartment, where no INDEL larger than 10kb was identified and our findings in the accessory compartment where few large INDELs were detected. Consequently, we find that the primary mode of sequence evolution in the core and accessory compartments occurs through the accumulation of single base pair mutations, as suggested by the negatively skewed distribution of SV-SNP index values (Fig. 4). In contrast, our pairwise comparisons reveal that *Cloud* sequences often differ from each other by hundreds of thousands of nucleotides that arise from multiple independently segregating INDELs greater than 10kb, and that on average, the amount of sequence impacted by these large INDELs vastly exceeds that impacted by SNPs. The mechanisms that promote large SV formation in *Starships* like *Cloud* remain unknown, although they are a focus of ongoing investigation. It is also intriguing that the diversifying effect of *Starships* is most prevalent in the clonally reproducing PoO strains. Future work is needed to determine if Starships are a major driver of genomic diversity in the absence of sexual reproduction. Importantly, by breaking down sequence-based analyses of *Starships* according to their family and navis classifications, our analysis takes an important step towards highlighting how *Starship* sequence evolution may deviate depending on both the identity of specific elements and the genomic background.

Another distinct and important contribution of *Starships* to genome diversification is their ability to transfer genetic information across species barriers. Although mini/accessory chromosomes similarly allow for the horizontal transfer of many genes simultaneously, the largest distance yet observed in fungi remains within genera (Habig et al. 2025) such as that of Metarhizium species that diverged ca. 15 million years ago (MYA) (Habig et al. 2024). In contrast, there is bioinformatic evidence of *Starship* transfer between different fungal orders that have diverged ca. 180 MYA, and both bioinformatic and laboratory-based evidence of transfer between genera diverging ca. 100 MYA (Urquhart et al. 2025). In *Verticillium* plant pathogens, there has been extensive horizontal transfer of *Starships* with other fungi and plant pathogens from different taxonomic orders, which possibly facilitated the acquisition of pathogenicity factors (Sato et al. 2025). *Starships* therefore increase the range of potential genetic donors and in turn expand the potential variety of novel genes that can be acquired horizontally by a given recipient lineage. Due to the complexity of mini chromosomes (high repeat content, complex rearrangements, variable presence) (Peng et al. 2019; Langner et al. 2021; Liu et al. 2024; Lin et al. 2025) we were unable to reliably detect segregating insertion sites associated with *Starship* elements. However, the presence of *Starship tyrR* coding sequences on mini-chromosomes raises the intriguing possibility that these two compartments interact or exchange material. Continued experimental evidence to determine if these mini-chromosome *tyrR* are active, and whether mini-chromosomes are the target of *Starship* insertions will be needed to answer this question. Although the phenotypic relevance of *Starship* transfers in *Pyricularia* remains to be elucidated, particularly as it regards the evolution of plant-fungal interactions, such transfers in other fungal species are often associated with the repeated emergence of adaptive phenotypes like formaldehyde resistance and even plant host-specific pathogenicity (A. S. Urquhart et al. 2024; Bucknell et al. 2025).

We extend the potential catalog of *Starship* phenotypes by identifying that well characterized effectors and avirulence coding sequences in *P. oryzae* are carried by *Starships* as genetic cargo. We further show that presence on a *Starship* appears to result in high levels of PAV, copy number variation, and sequence exchange, indicative of specific evolutionary consequences for cargo sequences located in the *Starship* compartment. Important host determinants such as Pwl1 and Pwl2 are both present on *Starships,* and their variable presence in *P. oryzae* strains can impact their detection by host plants and subsequent ability to infect (Sweigard et al. 1995; Masaki et al. 2023; Brabham et al. 2024). It was previously identified that *AvrPita1* and *AvrPita2* are present at multiple genomic locations across strains, while AvrPita3 was ubiquitous at a syntenic site between strains (Chuma et al. 2011). The *AvrPita1* and *AvrPita2* sequences are flanked by a retrotransposon that was proposed to mediate these multiple translocations, while the identification of *AvrPita1* on mini chromosomes prompted the hypothesis that mini-chromosomes serve as a genetic way station to maintain the sequence outside of core chromosomes and move between strains (Peng et al. 2019). Our detection of *AvrPita1* on a *Starship* now raises the possibility that *Starships* serve a similar mechanism to harbor and exchange coding sequences under diverging selective pressure (i.e. host detection versus virulence). We identified similar movement dynamics for the MAX07 coding sequence which is structurally similar to *AvrPiz-t*, indicating that multiple translocations across core chromosomes could be a recurring theme for *Starship* associated effectors. The detection of increased copy number for the *MoNudix-1* coding sequence and of putative BGC in association with a *Starship*, along with the detection of a duplication of an N-terminal amidases, all point to *Starships* being major drivers of sequence diversification that is positioned to impact the evolution of pathogen virulence and host detection.

There are many open questions regarding *Starship* regulation, maintenance, and transfer, many of which hinge upon how these elements are epigenetically regulated in fungal genomes. We provide a comprehensive epigenetic profile of *Starship* elements across multiple strains and find significant variation within and between families of elements that aligns with how genetic and activity variation is partitioned across these elements. The SLR and *Enterprise* family elements are on average more completely covered by both heterochromatic histone modifications, namely H3K27me3 and H3K9me3. Within the SLR elements, there is substantial variation for both marks between elements, particularly in strains O-135 and Br48. The functional consequence of this variation between elements remains an open question for further study. In contrast to the SLR and *Enterprise* family elements, *Arwing* family elements have significantly fewer heterochromatic histone modifications, which we suggest is directly associated with the fact that these elements show the greatest levels of intergenome PAV and evidence of horizontal transfer. All the 29 *Arwing* family elements identified in *P. oryzae* were classified as variable or unique. However, it is not universally true that all *Arwings* have a more euchromatic profile or that they are the only *Starship* family that shows evidence of mobility. We find that previously identified HGT events between *P. oryzae* (PoT) and *P. pennesetigina* (Kobayashi et al. 2023) in fact correspond to *Arwing*, *Enterprise*, and SLR elements, thus providing a mechanism capable of explaining the origins of horizontally acquired material in these fungal genomes. The horizontally transferred elements are generally covered by heterochromatic histone modifications in *P. oryzae* strain Br48, except for one *Arwing* element that has less than 20% coverage by H3K27me3 peaks. Likewise, while the majority of putative HGT events in the PoO strains are *Arwing* elements with low levels of H3K9me3 and H3K27me3, one *Arwing* in Guy11 has nearly 80% H3K27me3 coverage, indicating recent histone methylation and potential silencing in Guy11. Therefore, it appears that active genome defense via heterochromatin formation has successfully silenced *Starships* following more recent transfers. However, we note that *Arwing* family elements, and particularly *Arwings* in PoO strains, appear to be avoiding the accumulation of heterochromatic markers and that HGT of *Starships* does not per se trigger epigenetic silencing, given our finding of horizontally acquired elements without H3K27me3. An Arwing *Starship* similarly was found to mediate interstrain movement in *V. dahliae* and carried two well characterized effectors, *Ave1* and *Av2* (Sato et al. 2025). It is currently unclear whether these *Arwing* family elements are mobile because they have avoided genome silencing or if they have a mechanism for avoiding genome silencing and are therefore mobile.

Despite the acceptance and use of the core and accessory genome categories, it is often not clear what mechanisms drive their emergence and evolution. As such, there is a need to further improve genomic models to integrate biochemical and molecular features of the genome with evolutionary theory and data to provide a more complete understanding of the function and evolution of eukaryotic genomes. The core and accessory compartmentalization model provides one such approach, and its refinement via the inclusion of *Starships* stands to improve our ability to explain the emergence of genomic and epigenetic variation as well as ongoing patterns of gene transfer in nature (Urquhart et al. 2025; Gluck-Thaler et al. 2022; 2025).

## Figures

Figure 1. Diverse *Starship* and Starship-like regions (SLRs) characterize the *Starship* compartment of *P. oryzae*.

Figure 2. *Starships* in the Arwing family are highly variable across *P. oryzae* strains.

Figure 3. The *Starship* compartment of PoO strains contains the most variable regions of the genome.

Figure 4. C*loud Starships* contain multiple independent insertion, deletion, and duplication structural variants across strains and are hotspots for genomic variation.

Figure 5. Effector coding sequences in the *Starship* compartment have high interstrain variability and mobility.

Figure 6. *Cloud Starships* have undergone frequent horizontal transfer across the *Pyricularia* genus.

Figure 7. The Arwing family epigenome resembles that of the core compartment.

## Supplemental Figures

Fig. S1. Simplified schematic comparing the *Starship* annotation approaches of stargraph and starfish.

Fig. S2. Genome organization for all *Starship* and SLR elements identified in PoO strains

Fig. S3. Genome organization for all *Starship* and SLR elements identified in PoTLE strains

Fig. S4. Clustering of elements using the *k*-mer Jaccard similarity networks of the *Starship* compartment and accessory regions in *P. oryzae*.

Fig. S5. Variability in clusters built from genome compartments within the PoO lineage strains with the SLR and *Starship* elements split into separate networks.

Fig. S6. Comparison of *Starship* and SLR cluster variability between the *P. oryzae* strains and 13 *Aspergillus fumigatus* strains.

Fig. S7. Variation in cargo content within the *Cloud Starships* (Arwing family).

Fig. S8. Expanded copy number of Nudix effectors in PoO strains are associated with the *Starship* compartment.

Fig. S9. Evidence of horizontal transfer of SLR ‘EA18_SLR2’ to *P. grisea* (GCA002871045) using alignment and gene identity comparisons.

Fig. S10. Evidence of horizontal transfer of *Starship* ‘MoOO135_e00088’ to *P. pennisetigena* (GCA004337985) using alignment and gene identity comparisons.

Fig. S11. Evidence of horizontal transfer of *Starship* ‘MoOO137_s00097’ to *P. pennisetigena* (GCA003013045) using alignment and gene identity comparisons.

Fig. S12. Evidence of horizontal transfer of *Starship* ‘MZ5-1-6_s00130’ to *Magnaporthales sp.* (GCA003709005) using alignment and gene identity comparisons

Fig. S13. Evidence of horizontal transfer of *Starship* ‘PoP131_s00143’ to both *P. grisea* (GCA003933175) and *P. penniseti* (GCA041721605) using alignment and gene identity comparisons.

Fig. S14. Evidence of horizontal transfer of *Starship* ‘PoP131_s00144’ to two *P. grisea* genomes (GCA002871045 and GCA040113025) using alignment and gene identity comparisons.

Fig. S15. Epigenome peak coverage of *Starships* by strain and TE content of *Starships*.

Fig. S16. Previously described horizontal transfer events between *P. oryzae* and related *Pyricularia* species correspond to identified *Starships*.

Fig. S17. Genome-wide and compartment-specific RIP profiles in PoO and PoTLE strains.

## Supplemental Tables

Table S1. Publicly available *P. oryzae* genomes analyzed

Table S2. *Starship* and SLR information for all identified elements in 13 *P. oryzae* genomes

Table S3. Counts of family-navis *Starships* and SLRs in 13 *P. oryzae* strains

Table S4. Gene density and relative genome size of each genome compartment per strain

Table S5. Summary number and genome size of the *Starship* Compartment in 13 *P. oryzae* assemblies

Table S6. Description and stats of clusters extracted from a *k*-mer Jaccard similarity networks built from the Representative Core regions, Accessory regions and the *Starship* compartment (*Starships*+SLRs) in strains from the PoO lineage

Table S7. Description and stats of clusters extracted from a *k*-mer Jaccard similarity networks built from the *Starships* and SLRs separately in strains from the PoO lineage

Table S8. Description and stats of clusters extracted from a *k*-mer Jaccard similarity networks built from the Representative Core regions, Accessory regions and the *Starship* compartment (*Starships*+SLRs) in strains from the PoTLE lineage

Table S9. Description and stats of clusters extracted from a *k*-mer Jaccard similarity networks built from the *Starships* and SLRs separately in strains from the 13 *A. fumigatus* strains

Table S10. Average Nucleotide Identity (ANI) values from pairwise comparisons between long-read assemblies from within the PoO lineage, the PoTLE lineage or 13 *A. fumigatus* strains

Table S11. Sum of basepairs impacted by SNPs and INDELS called during pairwise alignment of different regions within compartment-specific network derived clusters

Table S12. Representative sequence for avirulence and infection characterized genes from *P. oryzae*.

Table S13. Representative sequence for the 94 MAX families identified in *P. oryzae*.

Table S14. Putative horizontal transfers of regions within the *Starship* compartment from *P. oryzae*

## METHODS

### De-novo annotations of assemblies

All *P. oryzae* assemblies (Table S1) used in our analysis were de-novo annotated using Earl Grey (Baril et al. 2024) for repetitive DNA and braker3 (Gabriel et al. 2024) for gene prediction (*--fungus*) using protein data from fungal BUSCOs (https://bioinf.uni-greifswald.de/bioinf/partitioned_odb11/Fungi.fa.gz) (*--prot_seq Fungi.fa.gz*) and compleasm (Huang and Li 2023) (*--busco_lineage sordariomycetes_odb10*). Genes were functionally annotated using InterProScan (Jones et al. 2014) (interproscan-5.67-99.0), EggNOG-mapper (Cantalapiedra et al. 2021) and funannotate (Palmer and Stajich 2020) set up using a funannotate database with the ascomycota lineage (*funannotate setup -b ascomycota*).

### *Starship* and SLR annotation

We developed a series of existing scripts for pangenome variation graph-based *Starship* detection (O’Donnell et al. 2025b) into a new open-access, user-friendly tool called stargraph (available from https://github.com/SAMtoBAM/stargraph). Stargraph functions by first generating a reference-free pangenome variation graph using pggb (Garrison et al. 2024) with the large segment length parameter set high in order to focus on large syntenic alignment blocks and reduced TE noise (-s 20000). Similarly, we set the haplotype/genome number (-n) which determines a maximum number of mappings to a query position, to the number of input assemblies, to discourage repetitive mapping within conserved regions. Following this, the genomes are broken into 1kb windows and presence-absence variants (PAVs) are extracted using odgi pav (Guarracino et al. 2022). This provides pair-wise evidence of coverage (1 to 1) for each 1kb window. Overlapping windows with less than 50% covered are merged with a maximum 5bp gap, windows smaller than 10kb are removed (filter out isolated TEs) and then the remaining regions are merged with a more permissive 20kb gap. Finally, all remaining regions below 30kb are removed, and the remaining >30kb regions are considered PAVs. PAVs are then annotated as *Starship*-related regions (SLRs) if they contain at least one *Starship-*specific gene, such as the tyrosine recombinase (DUF3435) or DUF3723 genes. Here we conservatively used only the *tyrR*.

We used stargraph (v1.0.0-beta) (default settings) (SAMtoBAM 2025) in conjunction with the previously published *Starship* annotation tool, starfish (Gluck-Thaler and Vogan 2024) for a comprehensive annotation of *Starships* and SLRs. Initially, all *Starships* were detected using both methods on 13 Long-read assemblies (Table S1). The coordinates from both methods were merged using *bedtools subtract*; keeping starfish calls preferentially (providing the bed file as -a) and removing all remaining split SLRs smaller than 30kb. We then manually evaluated all remaining calls using alignments and Jaccard similarity-based network analyses to see the remaining overlaps between the two different methods. In some cases, starfish’s extension elements (_e*), which are extended by aligning to a reference *Starship* sequence using BLAST, had their downstream border further extended to fit an overlapping SLR, i.e. the extension element overlapped with an SLR region prior to the subtraction step, but was too short based on alignments and therefore the downstream border was extended using the larger SLR coordinates that more appropriately aligned with other haplotypes. Lastly, some SLRs were reduced in size or removed. These SLRs were only detected due to a *Starship* inserting themselves into regions already containing presence/absence variation and therefore the PAV region was aggregated into a single SLR. We sorted all predicted *Starships* into navis and haplotypes groups based on *tyrR* captain orthology and pairwise *k*-mer sequence similarity, respectively, as previously described and implemented in the starfish pipeline (Gluck-Thaler and Vogan 2024). Representatives of each *Starship* haplotype were manually selected after visualizing all pairwise nucleotide alignments between elements using *nucmer* (Marçais et al. 2018) (*--maxmatch*) and gggenomes (Hackl et al. 2024).

### Accessory and Representative Core regions

The same set of accessory and representative core regions were defined once and maintained throughout all analyses. Accessory regions were defined as all PAVs detected in the pangenome variation graph produced by stargraph that did not contain any *tyrR* sequences (excluding mini chromosomes; n=667 accessory regions; mean, median length=77kb; SD=149kb). To represent the core compartment, we extracted a subset of regions consisting of +/-50kb surrounding n=50 randomly selected BUSCOs orthologs present in all genomes. These 50 representative regions contain a minimum of one BUSCO (*--busco_lineage sordariomycetes_odb10*) found as a single copy in all strains, and both the average lengths (median length=100kb) and total sum of bases (∼65Mb) are comparable to the *Starship* and accessory compartments (Fig. S3B; *Starships*/SLRs median length=142kb; Accessory median length=77kb; *Starships*/SLRs sum length=38Mb; Accessory sum length=86Mb).

### *Starship*/SLR *k*-mer network analyses

All *Starships* and SLRs were initially hard-masked using the Earl Grey-derived repeat annotation libraries (Baril et al. 2024), then had individual 31-mer *k*-mer ‘sketches’ built using sourmash (Irber et al. 2024) (4.9.0) (*sourmash sketch dna -p k=31,noabund –singleton)*. The sketches were then compared all vs all using both Jaccard similarity and max-containment values (*sourmash compare -k 31; sourmash compare -k 31 --max-containment*). Although Jaccard similarities provide a balanced and symmetric measure of shared *k*-mers, this statistic does not provide an intuitive representation of similarity between features of very different sizes. This is important to consider, as *Starships* and SLRs have a wide range in length, and due to the possibility of both SLRs being fragmented and nested insertions. The max-containment similarity is better at capturing these contained sequence similarities, but max-containment can also inflate the similarity due to the asymmetric comparisons. We therefore used the Jaccard similarity measure in our downstream network analyses in order to more conservatively define element clusters. Following *sourmash compare*, the pairwise similarity values were extracted and similarities below 10% were removed. In the network, clusters were simply defined as any group of nodes with any connecting edges. The control datasets of accessory and representative core regions were analysed in the same fashion.

The *Aspergillus fumigatus Starship* and SLR networks were created using stargraph on 13 long-read assemblies *A. fumigatus* assemblies (GCA024220425, GCA028752175, GCA028752205, GCA032627265, GCA040126065, GCA040142845, GCA040142855, GCA040142875, GCA040142885, GCA040142945, GCA040142975, GCA949125545, SAMN28487501) and *k*-mer networks built using the same parameters. All network plots are generated using ggraph (Pedersen 2025) with Fruchterman Reingold layouts (*layout = ‘fr’*). Jaccard similarity networks were used to define clusters as all edge-connected nodes.

### Variant calling between regions in network clusters

The two largest clusters defined in the Jaccard similarity network were extracted and pairwise aligned to one another using Minimap2 (*-cx asm20 --cs --secondary=no*) (Li 2018). Variants (SNPs and INDELs of various lengths) were called using paftools (*paftools.js call -L 50 −l 50*). Since paftools does not genotype large INDELs, we developed a custom script to parse the paf alignments to identify large INDELs (see ‘Data Availability’). These variants were identified as gaps between two consecutive alignments >10kb in size, thereby conservatively calling variants only flanked by alignments at both edges and not derived from missing alignments at the terminal ends of the regions. The same parameters were used in the two largest accessory network clusters and all clusters within the network of representative core regions.

### Horizontal Gene Transfer (HGT) analysis

All *Starships* and SLRs were analyzed for evidence of HGT using nucleotide-based sequence similarity searches with BLAST against a database of 10,010 *Pezizomycotina* genomes downloaded from NCBI (all genomes publicly available as of 20th December 2024, removing metagenomic assemblies and those flagged as ‘atypical; *datasets download genome taxon pezizomycotina --filename pezizomycotina_ncbi.zip --exclude-atypical --dehydrated --assembly-source genbank --mag exclude*). After initially filtering out individually aligned blocks less than 10kb and 80% identity, any remaining genome was removed if the total length of the alignment per *Starship*/SLR haplotype query was less than 50kb or covered <50% of the total query length. Each alignment was then realigned with nucmer (Marçais et al. 2018) with an additional 50kb added to the distal edges of the most upstream and downstream BLAST alignments to specifically visualize alignments between the putatively transferred region and not the surrounding content. The delta file was then converted to paf format using paftools delta2paf (Li 2018) and visualized using gggenomes (Hackl et al. 2024). For each pairwise genome comparison with an alignment-supported HGT event, we sought genome-wide evidence of HGT by generating ‘BLASTall’ plots (e.g. Figure 6D) based on an all vs all BLASTn search of all predicted gene transcripts in the *Starship*-containing query genome vs all nucleotide sequences in the target genome (*blastn -dust no -max_target_seqs 1 -max_hsps 1*). We visualized hits based on whether they were found within the *Starship* of interest or not. In the case of Br48_s00012 used in Figure 6D, we further grouped hits found outside of the *Starship* into accessory regions, representative core regions or outside of any defined compartments. All alignments and BLASTall plots were manually inspected and verified.

Eleven genomes belonging to four different *Pyricularia* species (GCA003933175, GCA041721605, GCA003709005, GCA002871045, GCA004337975, GCA944989265, GCA019328425, GCA944952825, GCA003013045, GCA004337985, GCA040113025) were identified as HGT donor/recipient candidates based on the nucmer and BLASTall alignments. These eleven genomes, plus two *Gaeumannomyces* genomes as outgroups (downloaded from https://doi.org/10.5281/zenodo.14823851 (Hill et al. 2025)) were combined with the 13 long-read *P. oryzae* assemblies to determine their phylogenetic relationships by building a BUSCO-based maximum likelihood species tree. We used the species tree to validate the NCBI species ID, which included *P. grisea*, *P. pennisetigena*, *P. penniseti*, ‘*Pyricularia sp*.’ and ‘*Magnaporthales sp*.’. The species tree for all 26 genomes was built by extracting 3921 BUSCO protein sequences (*-l sordariomycetes*) conserved across all genomes. Sequences were aligned with mafft (*--auto*) and trimed by trimal (*-automated1*). We constructed a maximum likelihood species tree using IQ-TREE2 (*--alrt 1000 --ufboot 1000 -T AUTO --seed 111111*). The tree was rooted using the *Gaeumannomyces* genomes.

### Biosynthetic Gene Cluster characterization

BGCs present in *Starships*/SLRs were manually annotated using functional annotations from Interpro and Eggnog. Additionally, antiSMASH (v8) (Blin et al. 2025) was used to confirm and refine the BGC characterization and look for similarities to known clusters.

### Effector and Virulence coding sequence identification

The location of 94 MAX effector coding sequences (Le Naour—Vernet et al. 2023) and 22 reported virulence and avirulence effectors (referred to here as effectors for simplicity) (previously reported (Zhang et al. 2021) (Table S12) were determined using GMAP (Wu and Watanabe 2005) and BLAST+ (Camacho et al. 2009). For the MAX effectors, a single representative sequence from the cluster was used for the search (Table S13) as there is low sequence diversity in the clusters between strains. The GMAP output for each strain was visual inspected for duplicate loci being hit against similar MAX sequences because of the expanded orthogroups previously reported (Le Naour—Vernet et al. 2023). These loci were cross-referenced against the BLAST hit and a single MAX was assigned per loci for the best matching MAX. The MAX and effector sequences were overlapped with the *Starship* compartment using bedtools (Quinlan and Hall 2010) and categorized as on a *Starship* or not. Hits to mini-chromosomes were removed as *Starships* were not annotated on these elements. Counts of MAX coding sequences and effectors were summarized using custom python scripts and plotted using Complex Heatmap in R. The PAV for MAX families was summarized from previous analysis (Le Naour—Vernet et al. 2023) (Table S3). For each MAX group, the number of times the sequence was reported as “Missing” out of the 119 was used to calculate fraction absent. Each MAX or effector sequence identified in the *Starship* compartment in at least one strain was used to summarize counts in individual *Starship* or SLR elements and the core genome using bedtools. The location of MAX07 was identified from the GMAP and BLAST results and plotted using gggenomes (Hackl et al. 2024). The *MoNudix-1* and *–2* coding sequences (XP_003711159.1 and KAI7909397.1) (McCombe et al. 2025) were identified in each *P. oryzae* genome using default tblastn search and plotted using gggenomes.

### RIP analysis

Genome wide RIP profiles and Large RIP Affected Regions (LRARs) were calculated using the RIPper webserver with modified default parameters (Wyk et al. 2019) (*Window Size: 2000; maximum substrate: 1; minimum product: 0.85; Composite Index Chain: 3*). CSV files were downloaded for both genome-wide RIP profiles and LRARs. In order to calculate the proportion of LRARs covered by different compartments; we generated 1000 randomisations of each compartment (only SLRs, only *Starships*, both SLRs and *Starships*, accessory or core) using *bedtools shuffle* on their respective bed files. We compared the randomized results to the real dataset using a Wilcoxon test with FDR correction for multiple comparisons. Similarly, we calculated the proportion of each compartment covered by LRARs by generating similarly 1000 randomisations of the LRAR positions and compared this to the real dataset.

### Epigenome analysis

We previously generated chromatin immunoprecipitation sequencing (ChIP-seq) data for the PoO strains (Zhang et al. 2021; Rowe et al. 2023). ChIP-seq data for H3K27me3 and H3K9me3 in Br48 was downloaded from the DDBJ Sequence Read Archive (Kobayashi et al. 2023). Genomic enrichment of each histone modification was determined via peak calling using MACS2 (Zhang et al. 2008). Peaks were then intersected with representative core, accessory, *Starship,* and SLR annotations to calculate a percent coverage of every element. Kruskal-Wallis tests with post-hoc Dunn’s tests were used to test for statistical differences between the coverage of each category. TEs were annotated in all 13 genomes with RepeatMasker (Smit et al. 2015) using a TE library generated from representative genomes of each pathotype (Nakamoto et al. 2023). The subsequent TE annotations were used to calculate the percentage of each *Starship* and SLR made up of TE sequence.

## Data availability

All data used to generate figures, and in-house scripts are publicly available at this figshare repository (10.6084/m9.figshare.30920822; https://figshare.com/s/b7df51fe8d126764d9d9). This includes a comprehensive Quarto R document with code for generating figures and analyses alongside the available inputs. The tool stargraph is available online at https://github.com/SAMtoBAM/stargraph and can be installed and run using both conda and singularity.

## Acknowledgements

We thank Aaron A. Vogan and Michael F. Seidl for helpful comments on an earlier draft of this manuscript. We thank Horacio D. Lopez-Nicora for helping initiate this collaborative work. We thank Annika Pratt and Jeanne Ropars for their help with troubleshooting stargraph.

## Supplementary Figures

**Figure S1.**
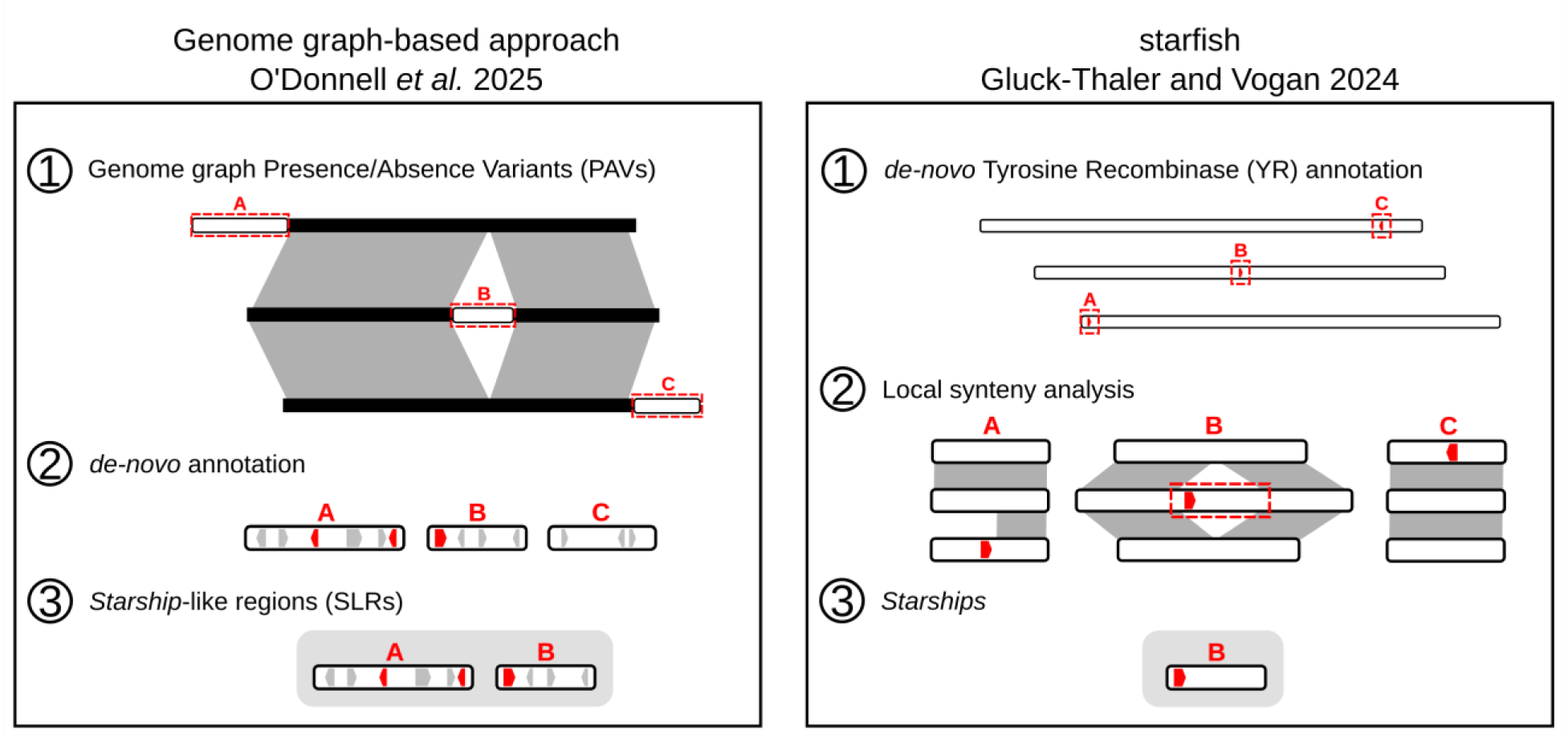
Simplified schematic comparing the *Starship* annotation approaches of stargraph and starfish. Two approaches were applied to annotating *Starships*. The first on the left uses a more permissive approach originally used in O’Donnell *et al*. 2025 and developed into a tool called stargraph. Stargraph uses a genome-graph approach, whereby (1) all genomes are aligned to one another and then Presence/Absence Variants (PAVs) are called based on all pairwise alignments in the graph. Following (2) gene annotation, (3) PAVs are considered *Starship*-like regions (SLRs) if they contain 1 or more *Starship*-specific genes such as the Tyrosine Recombinase (t*yrR).* The second approach on the right, using the tool starfish (Gluck-Thaler and Vogan 2024), is a more conservative that takes a tyrR forward approach. First, starfish (1) de-novo annotates all tyrRs then use their positions to (2) look for PAVs containing the gene. In this step, starfish is conservative in that it requires that (3) both edges of the PAV have alignments to a segregating site and that the *tyrR* is close to one edge of the inserted sequence.

**Figure S2.**
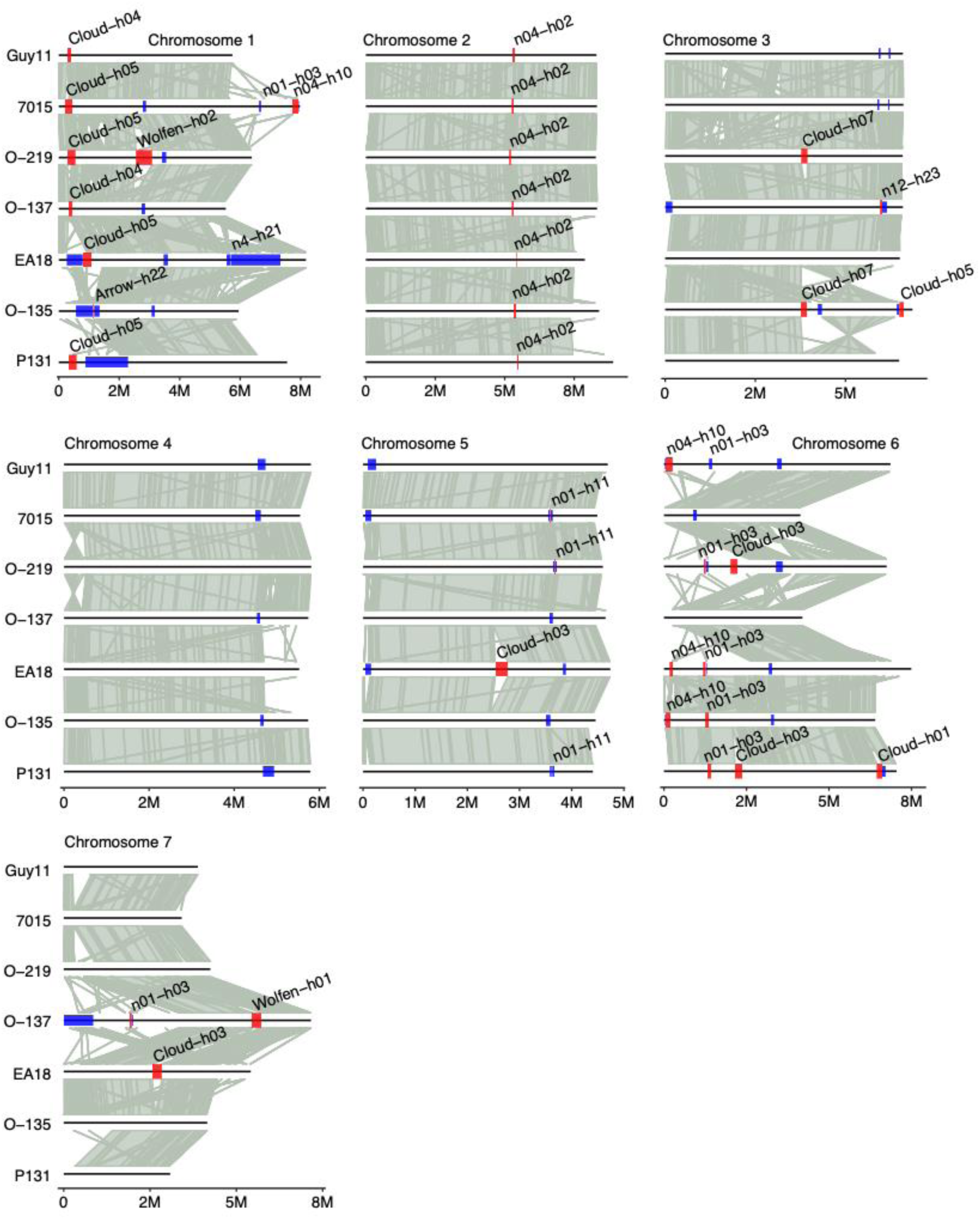
Genome organization for all *Starships* and SLRs identified in PoO strains. Chromosome synteny plots for the seven PoO strains. All seven chromosomes are labeled above each plot; strain names are labeled at the right, and chromosome lengths are labeled at the bottom in million base pairs (M). Grey links between chromosomes represent segments that are at least 90% sequence similar for at least 5 kb. *Starships* are shown as red boxes overtop the chromosome location and labeled with the navis-haplotype. *Starship-*like regions (SLRs) are shown as blue boxes.

**Figure S3.**
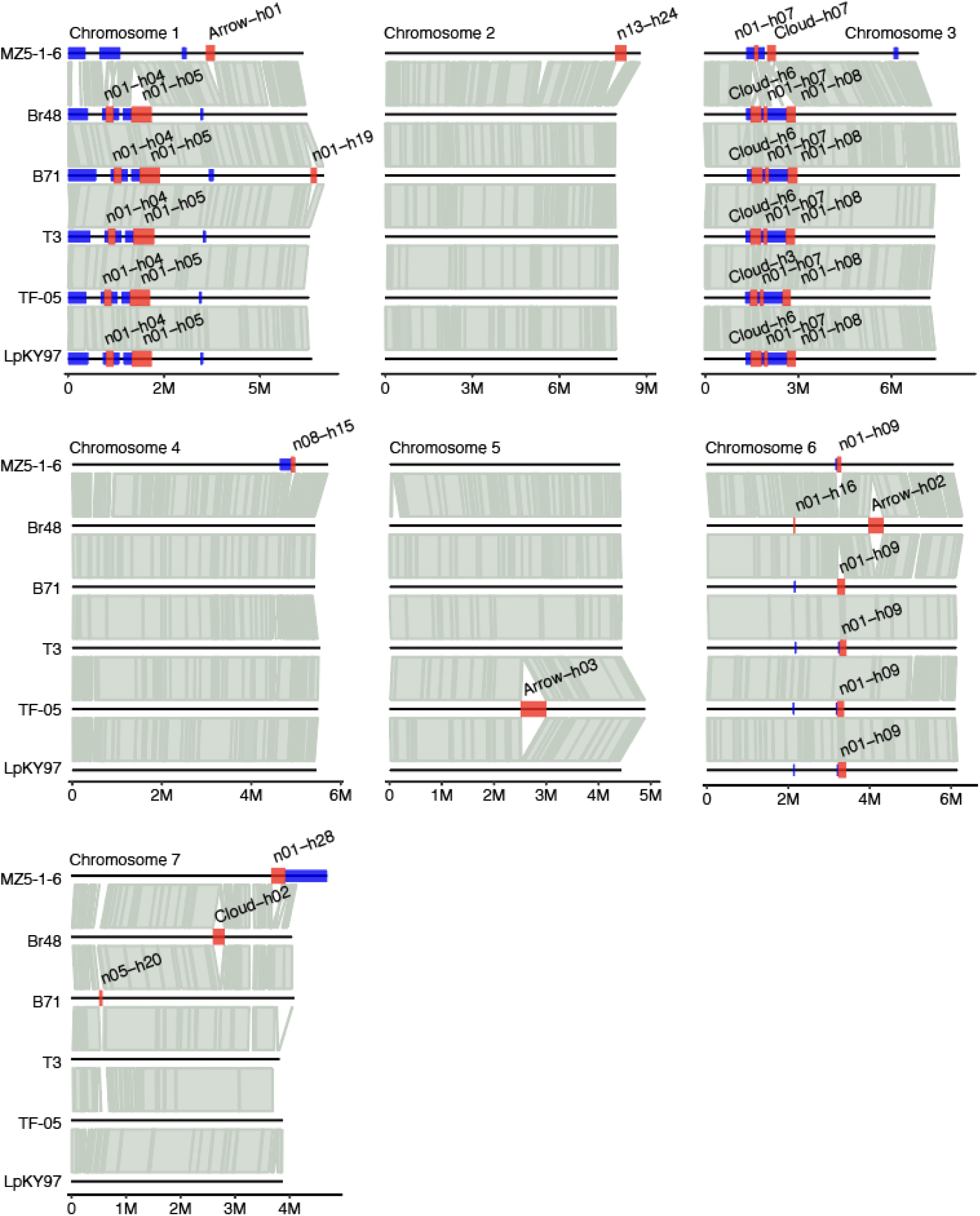
Genome organization for all *Starships* and SLRs identified in PoTLE strains. Chromosome synteny plots for the six PoTLE strains. All seven chromosomes are labeled above each plot; strain names are labeled at the right, and chromosome lengths are labeled at the bottom in million base pairs (M). Grey links between chromosomes represent segments that are at least 90% sequence similar for at least 5 kb. *Starships* are shown as red boxes overtop the chromosome location and labeled with the navis-haplotype. *Starship-*like regions (SLRs) are shown as blue boxes.

**Figure S4.**
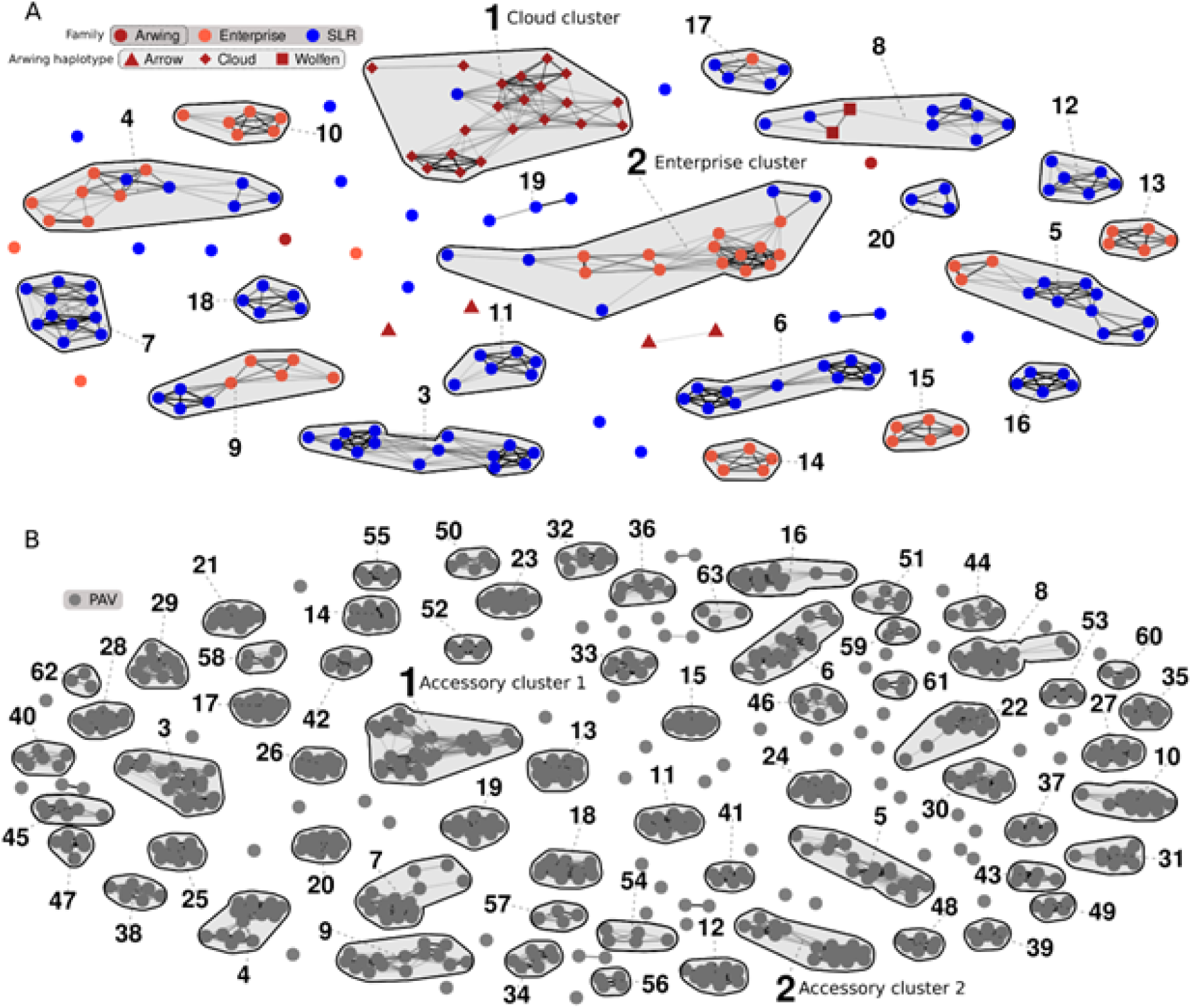
Clustering of elements using the *k*-mer Jaccard similarity networks of the *Starship* compartment and accessory regions in *P. oryzae*. Jaccard similarity networks were generated for the (A) *Starship* compartment and (B) accessory regions from all 13 *P. oryzae* genomes excluding mini-chromosomes. Clusters were determined as any set of ≥2 nodes connected by edges and are highlighted and numbered in order of largest to smallest based on the number of nodes (if an equal number of nodes, then the number of edges was used. If both nodes and edges were the same, the cluster number was assigned randomly). The larger numbered clusters 1 and 2 for both networks are those used to generate pairwise comparisons and call sequence variants.

**Figure S5.**
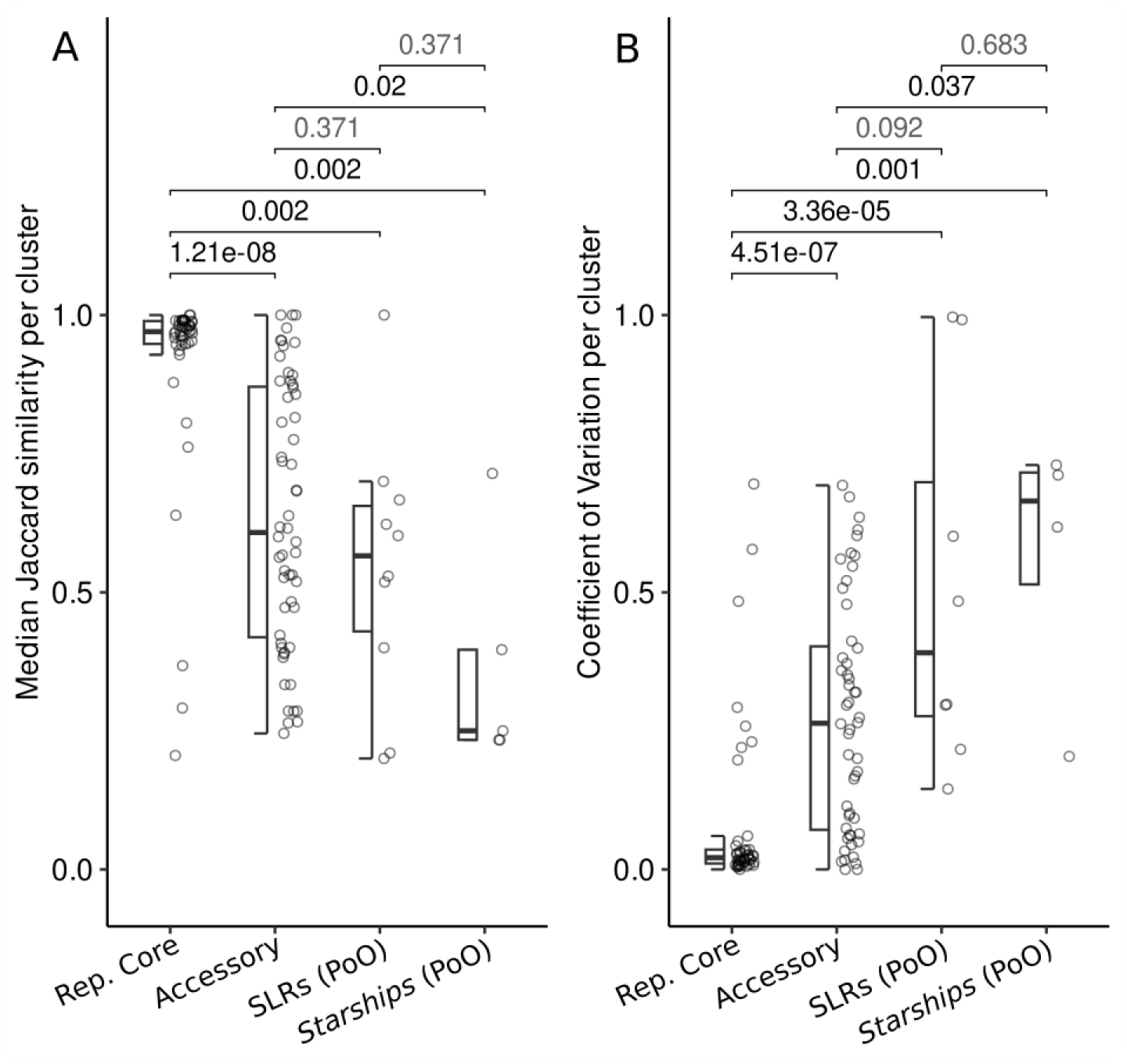
Variability in clusters built from genome compartments within the PoO lineage strains with the SLR and *Starship* elements split into separate networks. Jaccard similarity networks were separately built from regions of each defined compartment (Rep. Core, Accessory, SLRs and *Starships*) from strains from the PoO lineage. Clusters were defined as groups (n>2) of nodes connected by one or more edges. (A) Median Jaccard similarity within each cluster. (B) The coefficient of variation within each cluster. Wilcoxon test p-values with false discovery rate (FDR) corrections for multiple tests are shown above individual pairwise comparisons

**Figure S6.**
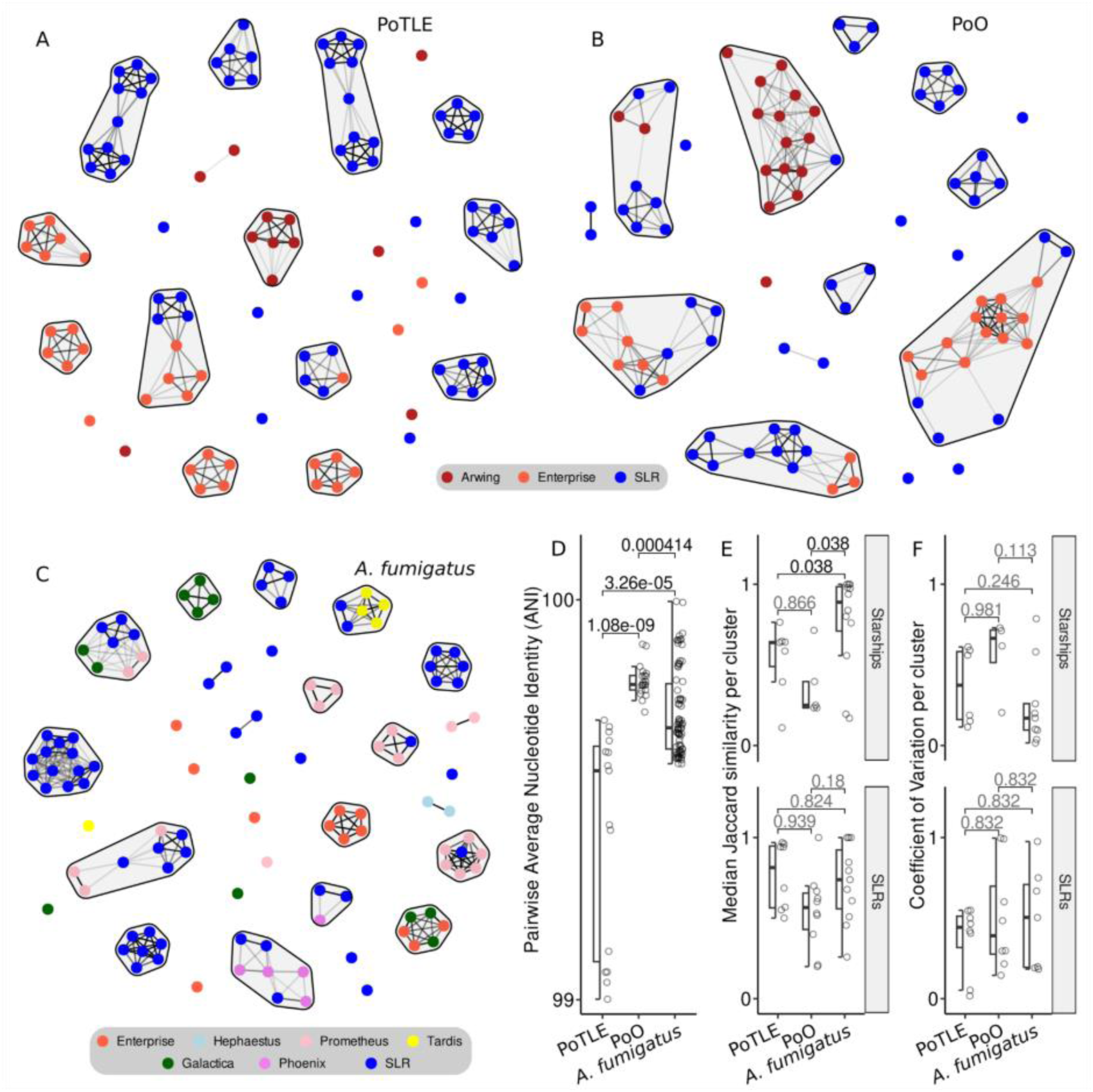
Comparison of *Starship* and SLR cluster variability between the *P. oryzae* strains and 13 *Aspergillus fumigatus* strains. The *Starship* compartment *k*-mer Jaccard similarity network is shown for the *P. oryzae* assemblies from the PoTLE lineage (A), the PoO lineage (B) and for 13 long-read *A. fumigatus* assemblies (C). *Starships* and SLRs are shown as nodes, edges are based on *k*-mer Jaccard similarity, and clusters are defined as any group of interconnected nodes. Node color represents the family of the *Starship tyrR* captain or SLR. (D) The *k*-mer based Average Nucleotide Identity (ANI) percentages from whole genome pairwise comparisons within each grouping. (E) The median Jaccard similarity and (F) coefficient of variation measured within each cluster defined in their respective networks and split between *Starships* and SLRs. Each cluster is depicted as open circles and summarized as box and whisker plots depicting the interquartile range and median shown as a thick black line. Wilcoxon test p-values are shown above individual pairwise comparisons with FDR correction for multiple comparisons.

**Figure S7.**
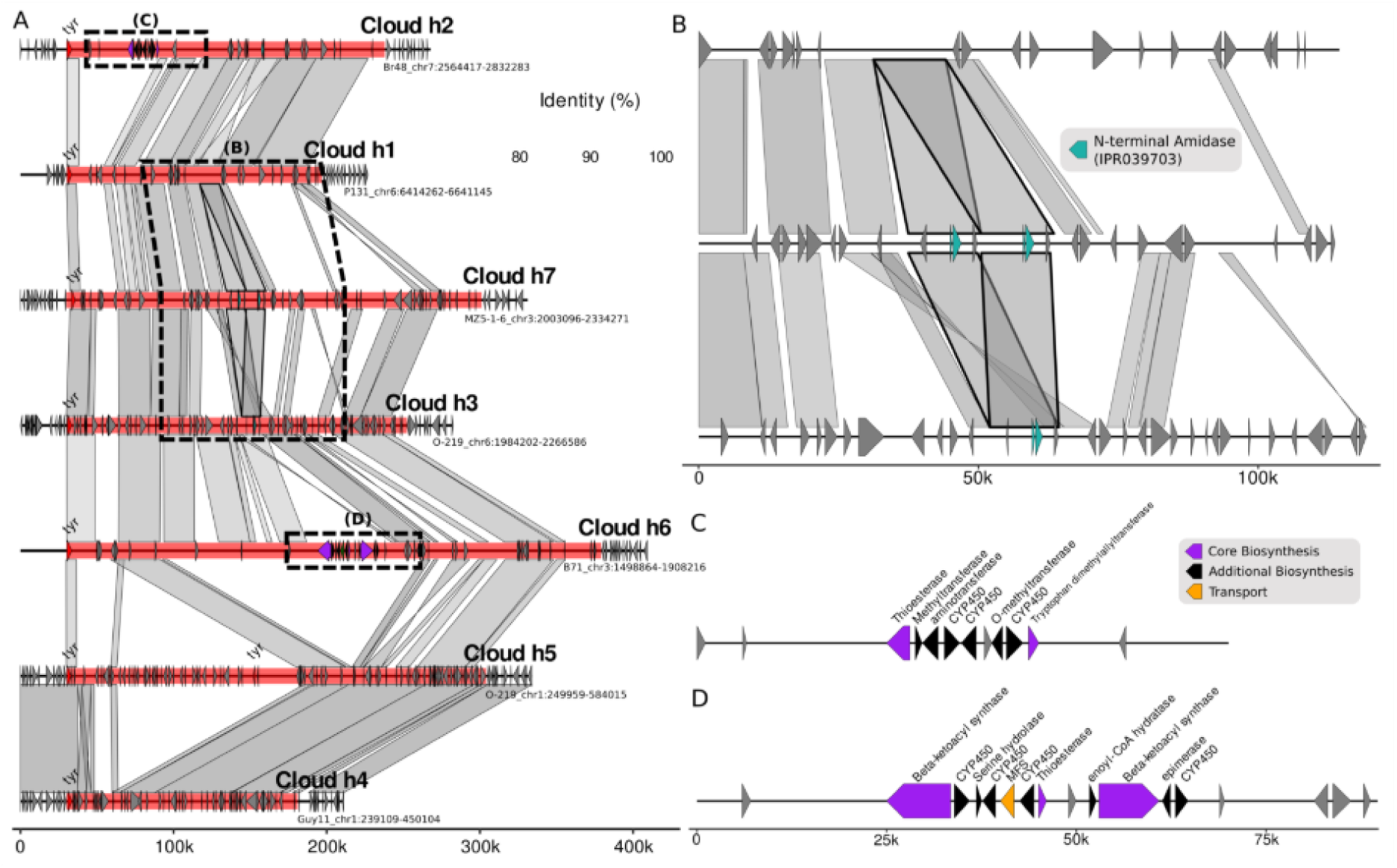
Variation in cargo content within the *Cloud Starships* (Arwing family). (A) Synteny plots of representatives from each of the *Cloud* haplotypes plus 25kb up and downstream, with insets corresponding to panels B, C and D outlined with dotted lines. (B) Highlighted region surrounding the N-terminal Amidase protein duplication in the representative sequence for *Cloud*-h7. (C-D) Putative Biosynthetic Gene Clusters (BGCs) found in *Cloud* haplotypes 2 and 6.

**Figure S8.**
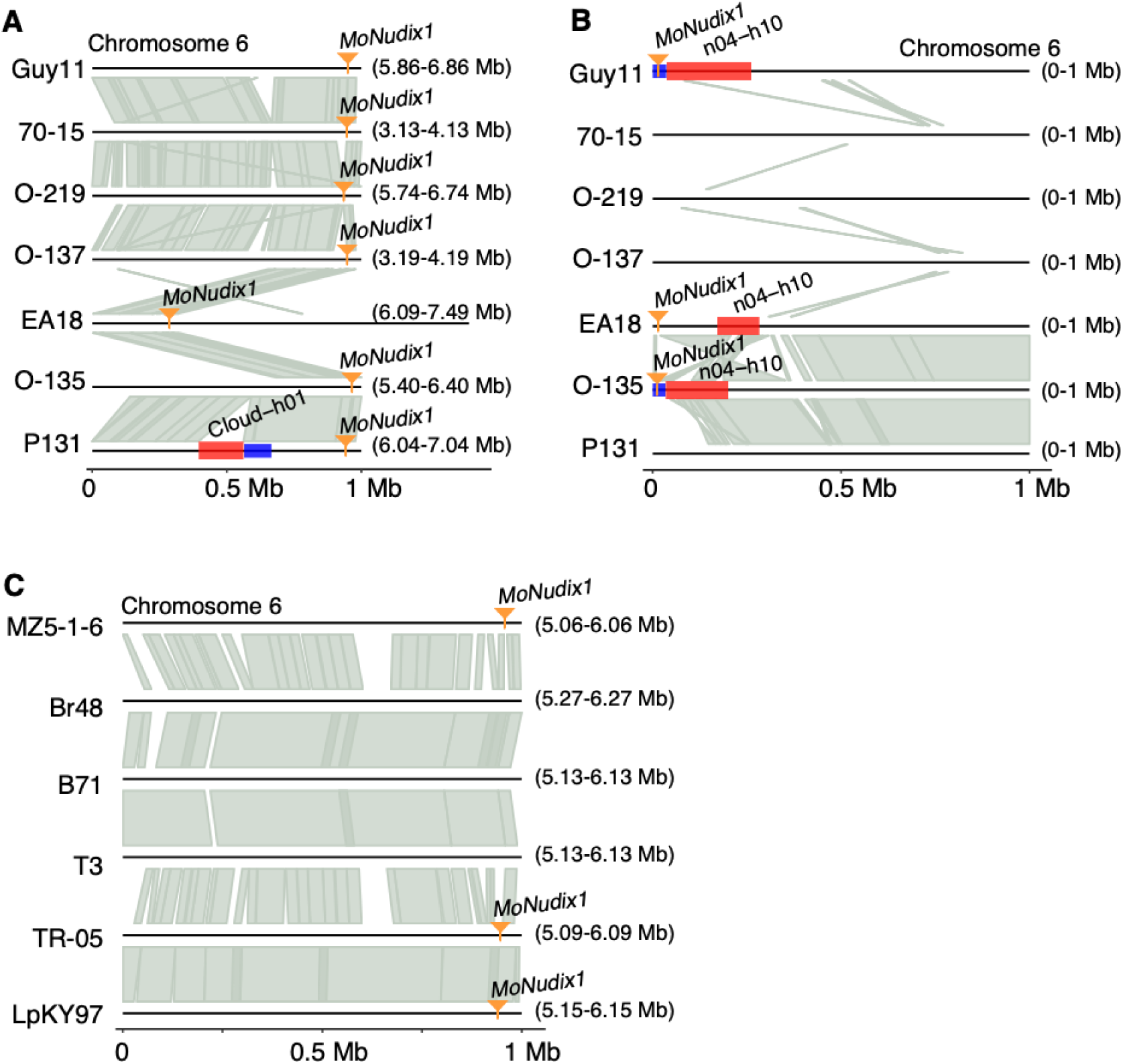
Expanded copy number of Nudix effectors in PoO strains are associated with the *Starship* compartment. Chromosome synteny plots around the *MoNUDIX-1* coding regions. Plot layout and details the same as described in Fig. S2. The chromosome location per strain is shown to the right (start-stop Mb). The position of the *MoNUDIX-1* coding sequence is indicated by an orange line and arrow. (A) The seven PoO strains all contain a copy of *MoNUDIX-1* at a similar region of chromosome 6. (B) Four PoO strains contain a second copy of *MoNUDIX-1* at the beginning of chromosome 6, which is in an SLR adjacent to a *Starship* in three strains. Note, the *MoNUDIX-1* SLR*-Starship* region is not depicted for strain 70-15 because there is a large chromosomal translocation in 70-15 from chromosome 6 to chromosome 1. The corresponding region can be seen in Fig. S2. (C) The one copy of *MoNUDIX-1* present in PoL and PoE strains is at a similar position to the shared copy in PoO strains.

**Figure S9.**
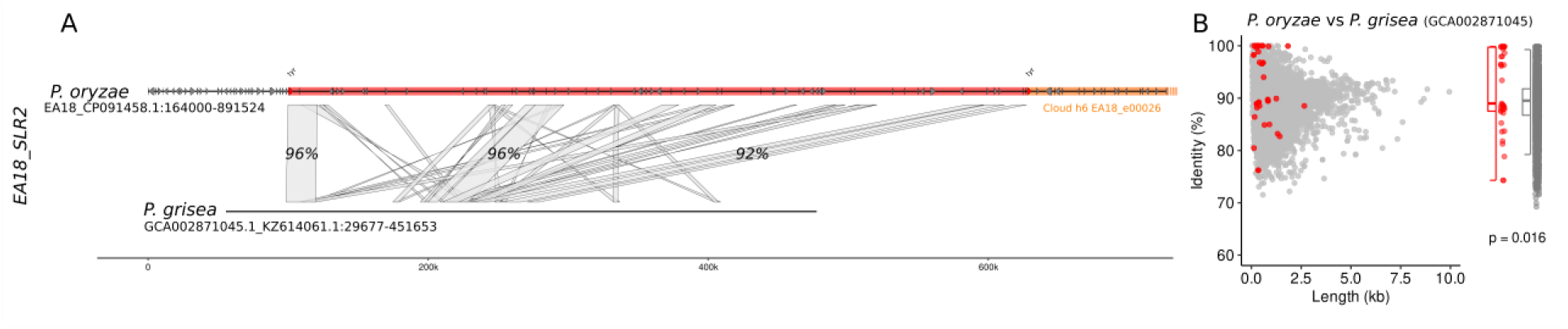
Evidence of horizontal transfer of SLR ‘EA18_SLR2’ to *P. grisea* (GCA002871045) using alignment and gene identity comparisons. (A) Alignment of EA18_SLR2 and *P. grisea* (GCA002871045) with a maximum of +/- 100kb either side. Highlighted region in red indicates the EA18_SLR2 element and *tyrR* genes are coloured in red and labelled. The orange coloured segment to the right of the EA18_SLR2 element is an adjacent *Starship* EA18_e00026 (*Cloud-*h6) that does not align the adjacent region in *P. grisea.* (B). Distribution of nucleotide identity and length values between all predicted gene sequences from strain EA18 and nucleotide sequences from *P. grisea* (GCA002871045). A Wilcoxon test was used to compare the difference in identity between genes within EA18_SLR2 (red) and all other genes (grey).

**Figure S10.**
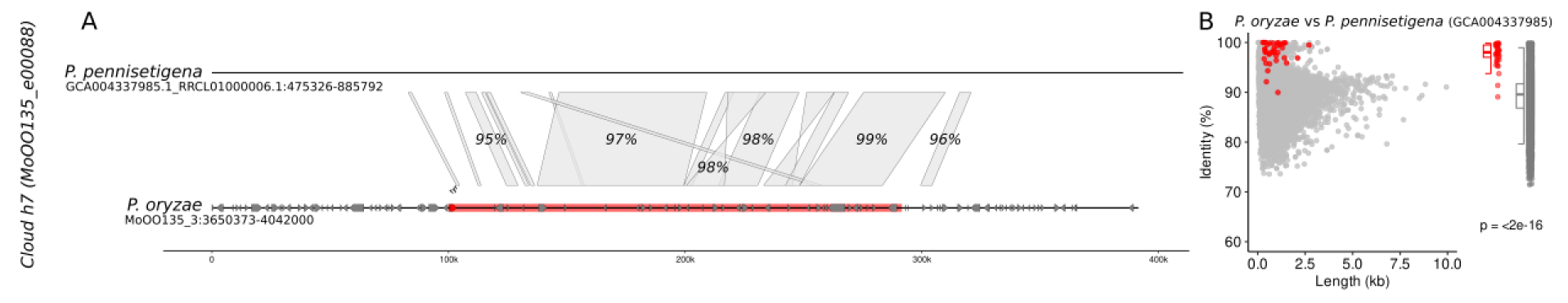
Evidence of horizontal transfer of *Starship* ‘MoOO135_e00088’ to *P. pennisetigena* (GCA004337985) using alignment and gene identity comparisons. (A) Alignment of MoOO135_e00088 and *P. pennisetigena* (GCA004337985) with a maximum of +/- 100kb either side. Highlighted region in red indicates the MoOO135_e00088 element and *tyrR* genes are coloured in red and labelled (B). Distribution of nucleotide identity and length values between all predicted gene sequences from strain O-135 and nucleotide sequences from *P. pennisetigena* (GCA004337985). A Wilcoxon test was used to compare the difference in identity between genes within MoOO135_e00088 (red) and all other genes (grey).

**Figure S11.**
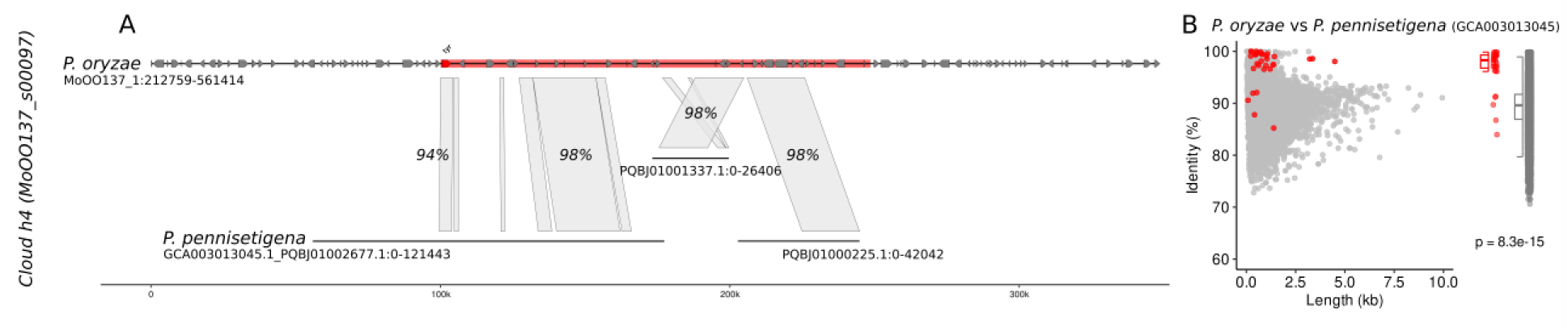
Evidence of horizontal transfer of *Starship* ‘MoOO137_s00097’ to *P. pennisetigena* (GCA003013045) using alignment and gene identity comparisons. (A) Alignment of MoOO137_s00097 and *P. pennisetigena* (GCA003013045) with a maximum of +/- 100kb either side. Highlighted region in red indicates the MoOO137_s00097 element and *tyrR* genes are coloured in red and labelled (B). Distribution of nucleotide identity and length values between all predicted gene sequences from strain O-137 and nucleotide sequences from *P. pennisetigena* (GCA003013045). A Wilcoxon test was used to compare the difference in identity between genes within MoOO137_s00097 (red) and all other genes (grey).

**Figure S12.**
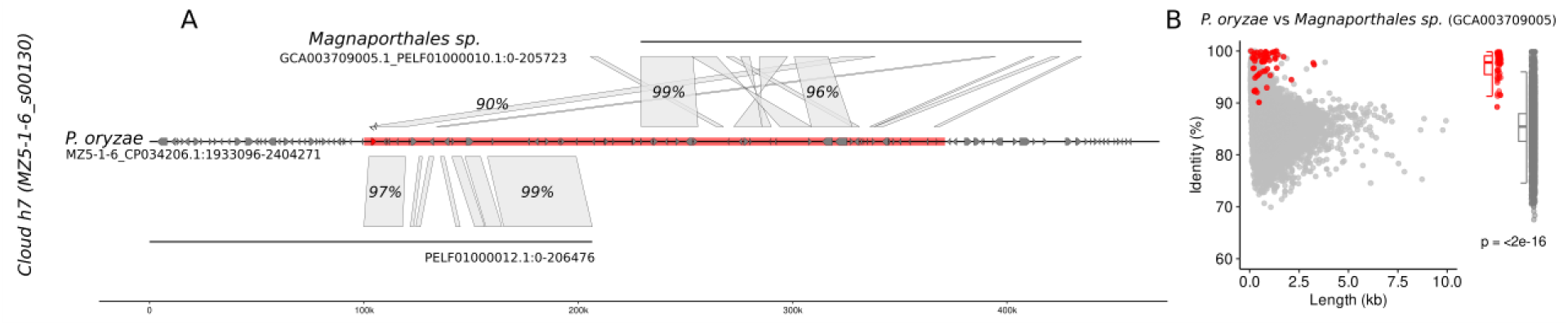
Evidence of horizontal transfer of *Starship* ‘MZ5-1-6_s00130’ to *Magnaporthales sp.* (GCA003709005) using alignment and gene identity comparisons. (A) Alignment of MZ5-1-6_s00130 and *Magnaporthales sp*. (GCA003709005) with a maximum of +/- 100kb either side. Highlighted region in red indicates the MZ5-1-6_s00130 element and *tyrR* genes are coloured in red and labelled (B). Distribution of nucleotide identity and length values between all predicted gene sequences from strain MZ5-1-6 and nucleotide sequences from *Magnaporthales sp*. (GCA003709005). A Wilcoxon test was used to compare the difference in identity between genes within MZ5-1-6_s00130 (red) and all other genes (grey).

**Figure S13.**
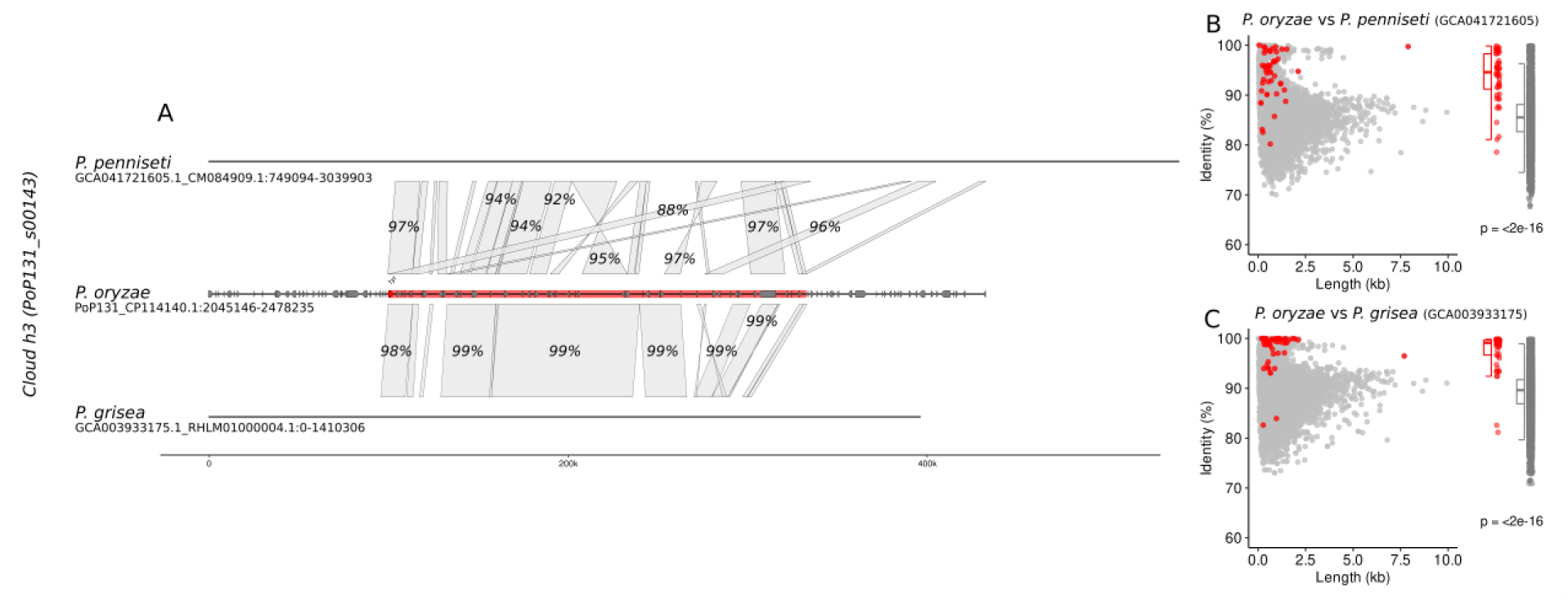
Evidence of horizontal transfer of *Starship* ‘PoP131_s00143’ to both *P. grisea* (GCA003933175) and *P. penniseti* (GCA041721605) using alignment and gene identity comparisons. (A) Alignment of PoP131_s00143 and *P. grisea* (GCA003933175) and *P. penniseti* (GCA041721605) with a maximum of +/- 100kb either side. Highlighted region in red indicates the PoP131_s00143 element and *tyrR* genes are coloured in red and labelled (B-C). Distribution of nucleotide identity and length values between all predicted gene sequences from strain P131 and nucleotide sequences separately from *P. grisea* (GCA003933175) and *P. penniseti* (GCA041721605). A Wilcoxon test was used to compare the difference in identity between genes within PoP131_s00143 (red) and all other genes (grey).

**Figure S14.**
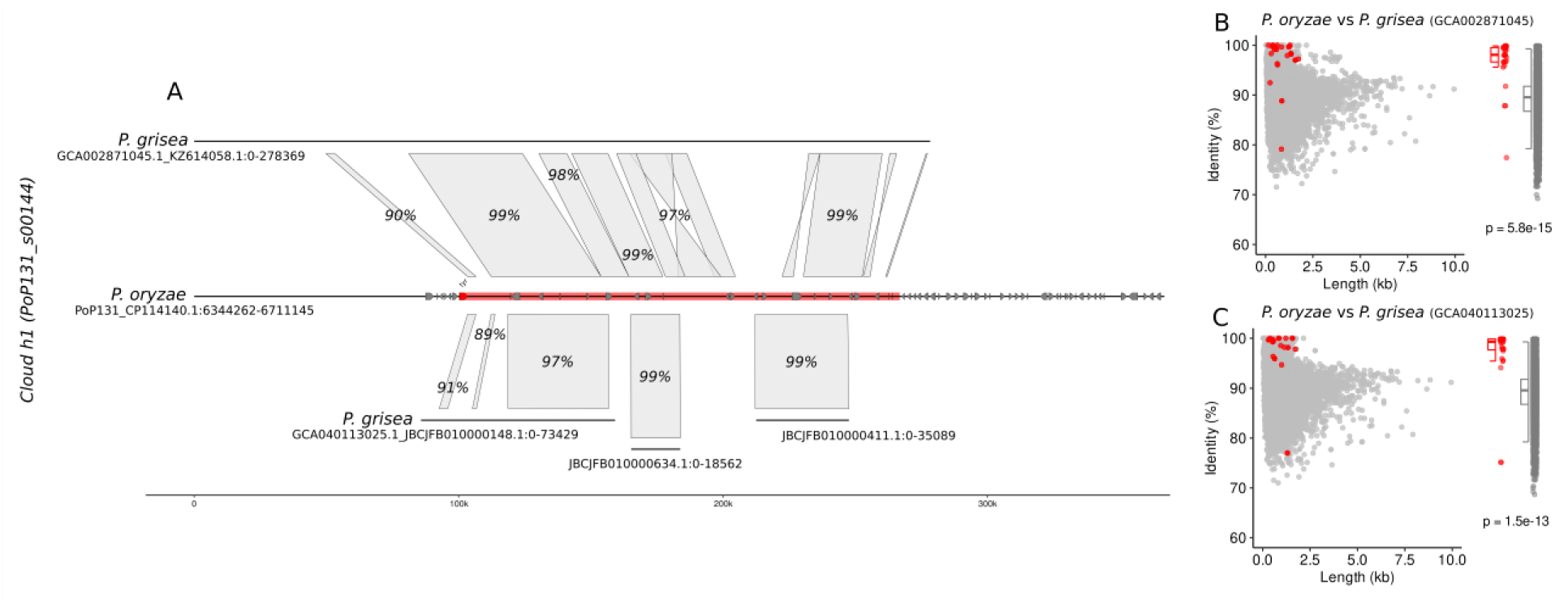
Evidence of horizontal transfer of *Starship* ‘PoP131_s00144’ to two *P. grisea* genomes (GCA002871045 and GCA040113025) using alignment and gene identity comparisons. (A) Alignment of PoP131_s00144 and two *P. grisea* genomes (GCA002871045 and GCA040113025) with a maximum of +/- 100kb either side. Highlighted region in red indicates the PoP131_s00144 element and *tyrR* genes are coloured in red and labelled (B-C). Distribution of nucleotide identity and length values between all predicted gene sequences from strain P131 and nucleotide sequences separately from *P. grisea* (GCA002871045 and GCA040113025). A Wilcoxon test was used to compare the difference in identity between genes within PoP131_s00144 (red) and all other genes (grey).

**Figure S15.**
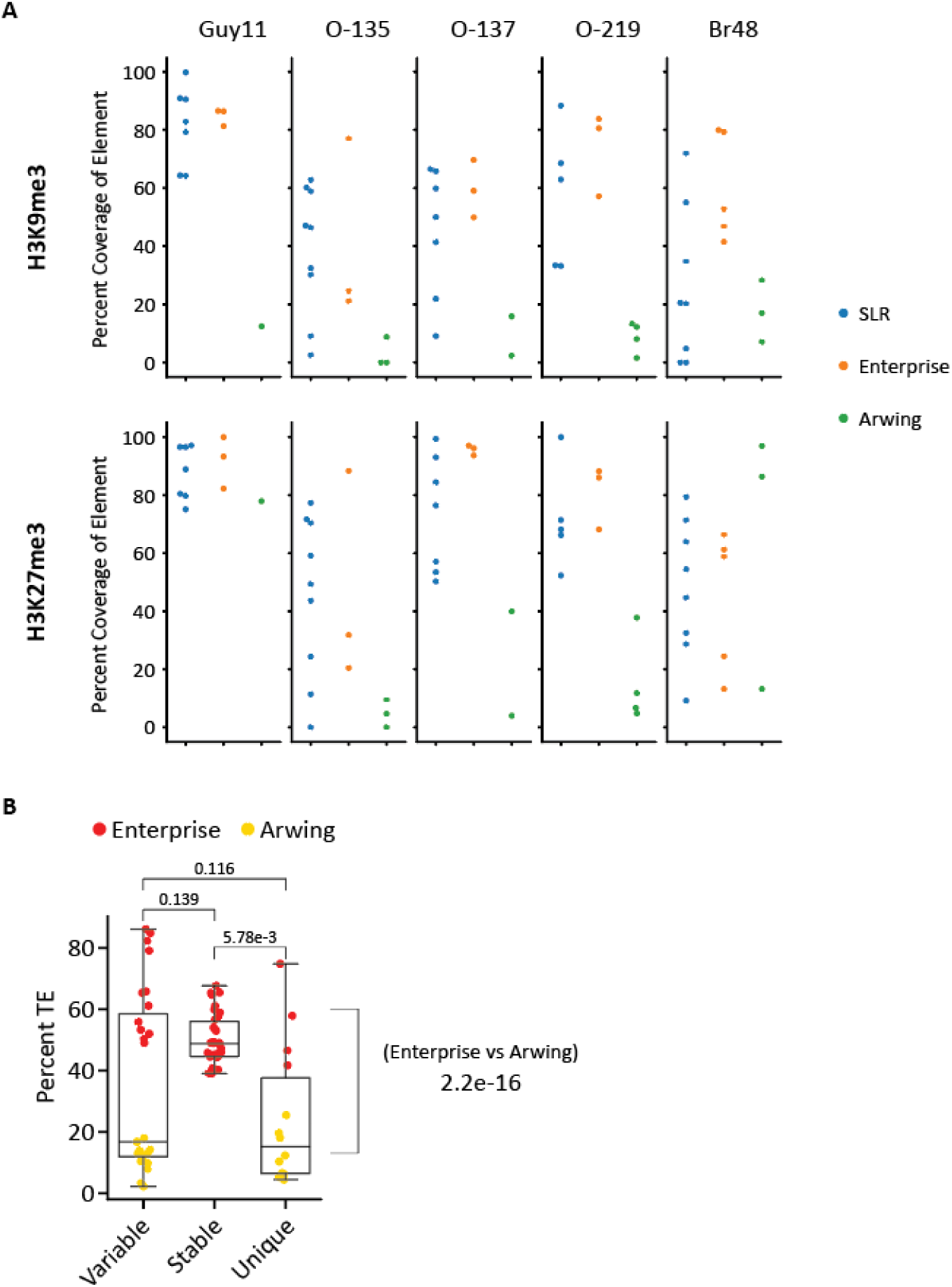
Epigenome peak coverage of *Starships* by strain and TE content of *Starships*. (A) Percent coverage of *Starships* and SLRs by H3K27me3 and H3K9me3 peaks as called by MACS2. Each strain is labeled at the top. Percent coverage is shown by element type-SLRs in blue, Enterprise *Starships* in orange, and Arwing *Starships* in green. (B) The percentage of each *Starship* made up of TEs. Each element is classified into variable, stable, or unique as described along the bottom, and further split between Enterprise (red) and Arwing (yellow) elements.

**Figure S16.**
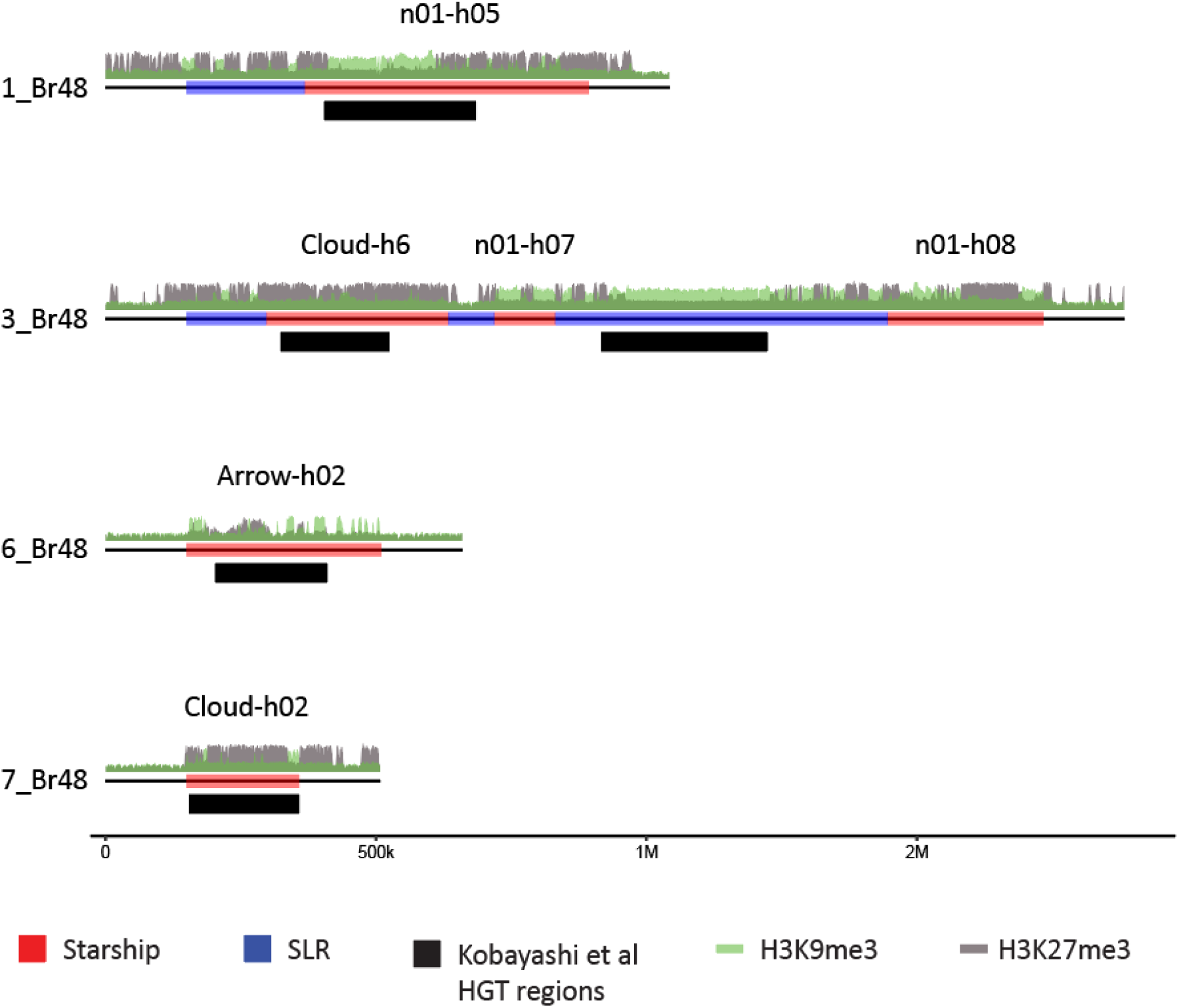
Previously described horizontal transfer events between *P. oryzae* and related *Pyricularia* species correspond to identified *Starships.*

**Figure S17.**
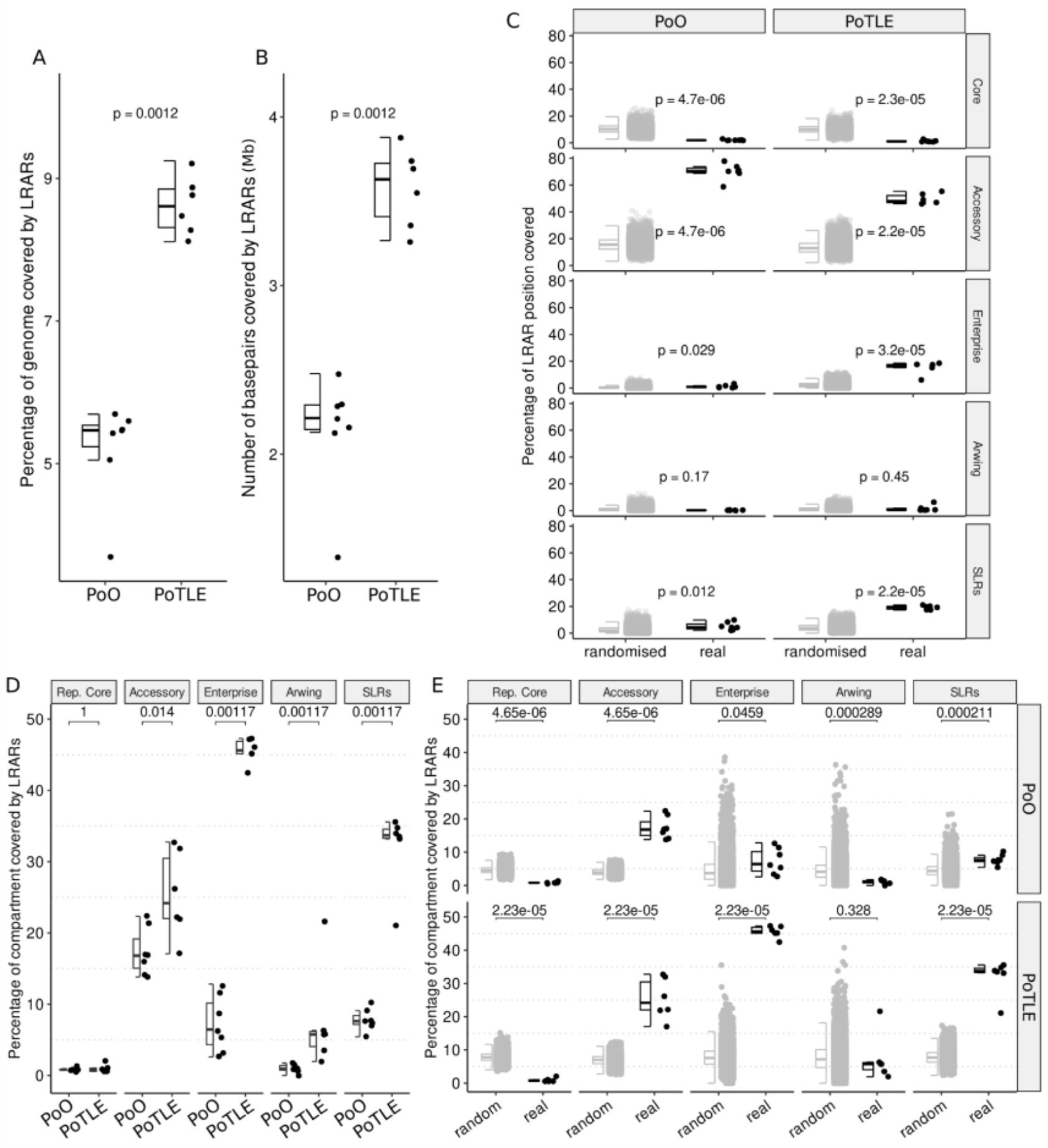
Genome-wide and compartment-specific RIP profiles in PoO and PoTLE strains. (A-B) Genome wide profile of Large RIP Associated Regions (LRARs) both as a percentage of genome size and total number of bases. (B) The total percentage of all LRARs detected in each genome covered by each genomic compartment (Rep. Core regions, Accessory regions, Enterprise elements, Arwing elements and *Starship-like* regions (SLRs) compared to a randomised distribution of LRARs. (D-E) The percentage of each compartment covered by LRARs were calculated using the genome wide LRAR pattern per strain. (D) Comparing LRAR coverage of each compartment between lineages. (E) Comparing the percentage of LRAR coverage between each compartment and a random distribution of LRARs split between each lineage. All p-values are from Wilcoxon tests with FDR corrections.

## REFERENCES

Allshire, Robin C., and Hiten D. Madhani. 2018. “Ten Principles of Heterochromatin Formation and Function.” Nature Reviews. Molecular Cell Biology 19 (4): 4. 10.1038/nrm.2017.119.

Badet, Thomas, Simone Fouché, Fanny E. Hartmann, Marcello Zala, and Daniel Croll. 2021. “Machine-Learning Predicts Genomic Determinants of Meiosis-Driven Structural Variation in a Eukaryotic Pathogen.” Nature Communications 12 (1): 3551. 10.1038/s41467-021-23862-x.

Baril, Tobias, James Galbraith, and Alex Hayward. 2024. “Earl Grey: A Fully Automated User-Friendly Transposable Element Annotation and Analysis Pipeline.” Molecular Biology and Evolution 41 (4): msae068. 10.1093/molbev/msae068.

Blin, Kai, Simon Shaw, Lisa Vader, et al. 2025. “antiSMASH 8.0: Extended Gene Cluster Detection Capabilities and Analyses of Chemistry, Enzymology, and Regulation.” Nucleic Acids Research 53 (W1): W32–38. 10.1093/nar/gkaf334.

Brabham, Helen J, Diana Gómez De La Cruz, Vincent Were, et al. 2024. “Barley MLA3 Recognizes the Host-Specificity Effector Pwl2 from Magnaporthe Oryzae.” The Plant Cell 36 (2): 447–70. 10.1093/plcell/koad266.

Bucknell, Angus, Hannah M. Wilson, Karen C. Gonçalves dos Santos, et al. 2025. “Sanctuary: A Starship Transposon Facilitating the Movement of the Virulence Factor ToxA in Fungal Wheat Pathogens.” mBio 16 (10): e01371–25. 10.1128/mbio.01371-25.

Bucknell, Angus, Hannah M. Wilson, Karen C. Gonçalves do Santos, et al. 2024. “Sanctuary: A Starship Transposon Facilitating the Movement of the Virulence Factor ToxA in Fungal Wheat Pathogens.” Preprint, bioRxiv, March 8. 10.1101/2024.03.04.583430.

Camacho, Christiam, George Coulouris, Vahram Avagyan, et al. 2009. “BLAST+: Architecture and Applications.” BMC Bioinformatics 10 (December): 421. 10.1186/1471-2105-10-421.

Cantalapiedra, Carlos P, Ana Hernández-Plaza, Ivica Letunic, Peer Bork, and Jaime Huerta-Cepas. 2021. “eggNOG-Mapper v2: Functional Annotation, Orthology Assignments, and Domain Prediction at the Metagenomic Scale.” Molecular Biology and Evolution 38 (12): 5825–29. 10.1093/molbev/msab293.

Chujo, Tetsuya, and Barry Scott. 2014. “Histone H3K9 and H3K27 Methylation Regulates Fungal Alkaloid Biosynthesis in a Fungal Endophyte-Plant Symbiosis.” Molecular Microbiology 92 (2): 2. 10.1111/mmi.12567.

Chuma, Izumi, Chihiro Isobe, Yuma Hotta, et al. 2011. “Multiple Translocation of the AVR-Pita Effector Gene among Chromosomes of the Rice Blast Fungus Magnaporthe Oryzae and Related Species.” PLoS Pathogens 7 (7): e1002147. 10.1371/journal.ppat.1002147.

Connolly, Lanelle R., Kristina M. Smith, and Michael Freitag. 2013. “The Fusarium Graminearum Histone H3 K27 Methyltransferase KMT6 Regulates Development and Expression of Secondary Metabolite Gene Clusters.” PLoS Genetics 9 (10): 10. 10.1371/journal.pgen.1003916.

Cook, David E, H Martin Kramer, David E Torres, Michael F Seidl, and Bart P H J Thomma. 2020. “A Unique Chromatin Profile Defines Adaptive Genomic Regions in a Fungal Plant Pathogen.” eLife 9 (December): e62208. 10.7554/eLife.62208.

Dhakal, Upasana, Hye-Seon Kim, and Christopher Toomajian. 2024. “The Landscape and Predicted Roles of Structural Variants in Fusarium Graminearum Genomes.” G3 Genes|Genomes|Genetics 14 (6): jkae065. 10.1093/g3journal/jkae065.

Dong, Suomeng, Sylvain Raffaele, and Sophien Kamoun. 2015. “The Two-Speed Genomes of Filamentous Pathogens: Waltz with Plants.” *Current Opinion in Genetics & Development*, Genomes and evolution, vol. 35 (December): 57–65. 10.1016/j.gde.2015.09.001.

Dong, Suomeng, and Yuanchao Wang. 2016. “Nudix Effectors: A Common Weapon in the Arsenal of Plant Pathogens.” PLoS Pathogens 12 (8): e1005704. 10.1371/journal.ppat.1005704.

Fang, Zhenyu, Yuyong Li, Jianqiang Huang, et al. 2025. “Experimental Insights into Genome Reconstruction Driven by Horizontal Transfer of Supernumerary Chromosomes in Magnaporthe Oryzae.” New Phytologist 248 (1): 140–56. 10.1111/nph.70438.

Fokkens, Like, Shermineh Shahi, Lanelle R. Connolly, et al. 2018. “The Multi-Speed Genome of Fusarium Oxysporum Reveals Association of Histone Modifications with Sequence Divergence and Footprints of Past Horizontal Chromosome Transfer Events.” Preprint, bioRxiv, November 7. 10.1101/465070.

Fouché, Simone, Ursula Oggenfuss, Emilie Chanclud, and Daniel Croll. 2022. “A Devil’s Bargain with Transposable Elements in Plant Pathogens.” Trends in Genetics 38 (3): 222–30. 10.1016/j.tig.2021.08.005.

Frantzeskakis, Lamprinos, Stefan Kusch, and Ralph Panstruga. 2019. “The Need for Speed: Compartmentalized Genome Evolution in Filamentous Phytopathogens.” Molecular Plant Pathology 20 (1): 1. 10.1111/mpp.12738.

Freitag, Michael. 2017. “Histone Methylation by SET Domain Proteins in Fungi.” Annual Review of Microbiology 71 (September): 413–39. 10.1146/annurev-micro-102215-095757.

Gabriel, Lars, Tomáš Brůna, Katharina J. Hoff, et al. 2024. “BRAKER3: Fully Automated Genome Annotation Using RNA-Seq and Protein Evidence with GeneMark-ETP, AUGUSTUS, and TSEBRA.” Genome Research 34 (5): 769–77. 10.1101/gr.278090.123.

Gardiner, Donald M., Megan C. McDonald, Lorenzo Covarelli, et al. 2012. “Comparative Pathogenomics Reveals Horizontally Acquired Novel Virulence Genes in Fungi Infecting Cereal Hosts.” PLOS Pathogens 8 (9): e1002952. 10.1371/journal.ppat.1002952.

Garrison, Erik, Andrea Guarracino, Simon Heumos, et al. 2024. “Building Pangenome Graphs.” Nature Methods 21 (11): 2008–12. 10.1038/s41592-024-02430-3.

Gladieux, Pierre, Bradford Condon, Sebastien Ravel, et al. 2018. “Gene Flow between Divergent Cereal- and Grass-Specific Lineages of the Rice Blast Fungus Magnaporthe Oryzae.” mBio 9 (1): 10.1128/mbio.01219-17. 10.1128/mbio.01219-17.

Gladieux, Pierre, Sébastien Ravel, Adrien Rieux, et al. 2018. “Coexistence of Multiple Endemic and Pandemic Lineages of the Rice Blast Pathogen.” mBio 9 (2): e01806–17. 10.1128/mBio.01806-17.

Gluck-Thaler, Emile, Adrian Forsythe, Charles Puerner, et al. 2025. “Giant Transposons Promote Strain Heterogeneity in a Major Fungal Pathogen.” mBio 16 (6): e01092–25. 10.1128/mbio.01092-25.

Gluck-Thaler, Emile, Timothy Ralston, Zachary Konkel, et al. 2022. “Giant Starship Elements Mobilize Accessory Genes in Fungal Genomes.” Molecular Biology and Evolution 39 (5): 5. 10.1093/molbev/msac109.

Gluck-Thaler, Emile, and Aaron A Vogan. 2024. “Systematic Identification of Cargo-Mobilizing Genetic Elements Reveals New Dimensions of Eukaryotic Diversity.” Nucleic Acids Research 52 (10): 10. 10.1093/nar/gkae327.

Gourlie, Ryan, Megan McDonald, Mohamed Hafez, et al. 2022. “The Pangenome of the Wheat Pathogen Pyrenophora Tritici-Repentis Reveals Novel Transposons Associated with Necrotrophic Effectors ToxA and ToxB.” BMC Biology 20 (1): 239. 10.1186/s12915-022-01433-w.

Gozashti, Landen, Daniel L. Hartl, and Russell Corbett-Detig. 2024. “Universal Signatures of Transposable Element Compartmentalization across Eukaryotic Genomes.” Preprint, bioRxiv, March 27. 10.1101/2023.10.17.562820.

Guarracino, Andrea, Simon Heumos, Sven Nahnsen, Pjotr Prins, and Erik Garrison. 2022. “ODGI: Understanding Pangenome Graphs.” Bioinformatics 38 (13): 3319–26. 10.1093/bioinformatics/btac308.

Habig, Michael, Anna V. Grasse, Judith Müller, Eva H. Stukenbrock, Hanna Leitner, and Sylvia Cremer. 2024. “Frequent Horizontal Chromosome Transfer between Asexual Fungal Insect Pathogens.” Proceedings of the National Academy of Sciences 121 (11): e2316284121. World. 10.1073/pnas.2316284121.

Habig, Michael, Cecile Lorrain, Alice Feurtey, Jovan Komluski, and Eva H. Stukenbrock. 2021. “Epigenetic Modifications Affect the Rate of Spontaneous Mutations in a Pathogenic Fungus.” Nature Communications 12 (1): 1. 10.1038/s41467-021-26108-y.

Habig, Michael, Satish Kumar Patneedi, Remco Stam, and Henrik Hjarvard De Fine Licht. 2025. “Horizontal Transfer of Accessory Chromosomes in Fungi – a Regulated Process for Exchange of Genetic Material?” Heredity, February 10, 1–7. 10.1038/s41437-025-00746-0.

Hackl, Thomas, Markus Ankenbrand, Bart van Adrichem, David Wilkins, and Kristina Haslinger. 2024. “Gggenomes: Effective and Versatile Visualizations for Comparative Genomics.” arXiv:2411.13556. Preprint, arXiv, November 5. 10.48550/arXiv.2411.13556.

Hill, Rowena, Michelle Grey, Mariano Olivera Fedi, et al. 2025. “Evolutionary Genomics Reveals Variation in Structure and Genetic Content Implicated in Virulence and Lifestyle in the Genus Gaeumannomyces.” BMC Genomics 26 (1): 239. 10.1186/s12864-025-11432-0.

Huang, Neng, and Heng Li. 2023. “Compleasm: A Faster and More Accurate Reimplementation of BUSCO.” Bioinformatics 39 (10): btad595. 10.1093/bioinformatics/btad595.

Irber, Luiz, N. Tessa Pierce-Ward, Mohamed Abuelanin, et al. 2024. “Sourmash v4: A Multitool to Quickly Search, Compare, and Analyze Genomic and Metagenomic Data Sets.” Journal of Open Source Software 9 (98): 6830. 10.21105/joss.06830.

Jones, Philip, David Binns, Hsin-Yu Chang, et al. 2014. “InterProScan 5: Genome-Scale Protein Function Classification.” Bioinformatics 30 (9): 1236–40. 10.1093/bioinformatics/btu031.

Kobayashi, Natsuki, Thach An Dang, Kieu Thi Minh Pham, et al. 2023. “Horizontally Transferred DNA in the Genome of the Fungus Pyricularia Oryzae Is Associated With Repressive Histone Modifications.” Molecular Biology and Evolution 40 (9): msad186. 10.1093/molbev/msad186.

Kramer, H. Martin, David E. Cook, Michael F. Seidl, and Bart P. H. J. Thomma. 2023. “Epigenetic Regulation of Nuclear Processes in Fungal Plant Pathogens.” PLOS Pathogens 19 (8): 8. 10.1371/journal.ppat.1011525.

Lahfa, Mounia, Philippe Barthe, Karine de Guillen, et al. 2024. “The Structural Landscape and Diversity of Pyricularia Oryzae MAX Effectors Revisited.” PLOS Pathogens 20 (5): e1012176. 10.1371/journal.ppat.1012176.

Langner, Thorsten, Adeline Harant, Luis B. Gomez-Luciano, et al. 2021. “Genomic Rearrangements Generate Hypervariable Mini-Chromosomes in Host-Specific Isolates of the Blast Fungus.” PLOS Genetics 17 (2): 2. 10.1371/journal.pgen.1009386.

Latorre, Sergio M., C. Sarai Reyes-Avila, Angus Malmgren, Joe Win, Sophien Kamoun, and Hernán A. Burbano. 2020. “Differential Loss of Effector Genes in Three Recently Expanded Pandemic Clonal Lineages of the Rice Blast Fungus.” BMC Biology 18 (1): 1. 10.1186/s12915-020-00818-z.

Le Naour—Vernet, Marie, Florian Charriat, Jérôme Gracy, et al. 2023. “Adaptive Evolution in Virulence Effectors of the Rice Blast Fungus Pyricularia Oryzae.” PLOS Pathogens 19 (9): e1011294. 10.1371/journal.ppat.1011294.

Li, Heng. 2018. “Minimap2: Pairwise Alignment for Nucleotide Sequences.” Bioinformatics 34 (18): 3094–100. 10.1093/bioinformatics/bty191.

Li, Wei, Baohua Wang, Jun Wu, et al. 2009. “The Magnaporthe Oryzae Avirulence Gene AvrPiz-t Encodes a Predicted Secreted Protein That Triggers the Immunity in Rice Mediated by the Blast Resistance Gene Piz-t.” Molecular Plant-Microbe Interactions: MPMI 22 (4): 4. 10.1094/MPMI-22-4-0411.

Lin, Guifang, Huakun Zheng, Dal-Hoe Koo, et al. 2025. “Highly Dynamic Supernumerary Mini-Chromosomes in a Magnaporthe Oryzae Strain Contributes to Cellular Variance of Genomic Content.” Fungal Genetics and Biology, 104020.

Liu, Sanzhen, Guifang Lin, Sowmya R. Ramachandran, et al. 2024. “Rapid Mini-Chromosome Divergence among Fungal Isolates Causing Wheat Blast Outbreaks in Bangladesh and Zambia.” The New Phytologist 241 (3): 3. 10.1111/nph.19402.

Marçais, Guillaume, Arthur L. Delcher, Adam M. Phillippy, Rachel Coston, Steven L. Salzberg, and Aleksey Zimin. 2018. “MUMmer4: A Fast and Versatile Genome Alignment System.” PLOS Computational Biology 14 (1): e1005944. 10.1371/journal.pcbi.1005944.

Masaki, Hosea Isanda, Santie de Villiers, Peng Qi, et al. 2023. “Host Specificity Controlled by PWL1 and PWL2 Effector Genes in the Finger Millet Blast Pathogen Magnaporthe Oryzae in Eastern Africa.” Molecular Plant-Microbe Interactions® 36 (9): 584–91. 10.1094/MPMI-01-23-0012-R.

McCombe, Carl L., Alex Wegner, Louisa Wirtz, et al. 2025. “Plant Pathogenic Fungi Hijack Phosphate Signaling with Conserved Enzymatic Effectors.” Science 387 (6737): 955–62. 10.1126/science.adl5764.

McNally, Alan, Yaara Oren, Darren Kelly, et al. 2016. “Combined Analysis of Variation in Core, Accessory and Regulatory Genome Regions Provides a Super-Resolution View into the Evolution of Bacterial Populations.” PLoS Genetics 12 (9): e1006280. 10.1371/journal.pgen.1006280.

Möller, Mareike, John B. Ridenour, Devin F. Wright, Faith A. Martin, and Michael Freitag. 2023. “H4K20me3 Is Important for Ash1-Mediated H3K36me3 and Transcriptional Silencing in Facultative Heterochromatin in a Fungal Pathogen.” PLOS Genetics 19 (9): 9. 10.1371/journal.pgen.1010945.

Möller, Mareike, Klaas Schotanus, Jessica L. Soyer, et al. 2019. “Destabilization of Chromosome Structure by Histone H3 Lysine 27 Methylation.” PLOS Genetics 15 (4): 4. 10.1371/journal.pgen.1008093.

Nakamoto, Anne A, Pierre M Joubert, and Ksenia V Krasileva. 2023. “Intraspecific Variation of Transposable Elements Reveals Differences in the Evolutionary History of Fungal Phytopathogen Pathotypes.” Genome Biology and Evolution 15 (12): evad206. 10.1093/gbe/evad206.

O’Donnell, Samuel, Gabriela Rezende, Jean-Philippe Vernadet, Alodie Snirc, and Jeanne Ropars. 2025a. “Harboring Starships: The Accumulation of Large Horizontal Gene Transfers in Domesticated and Pathogenic Fungi.” Genome Biology and Evolution 17 (7): evaf125. 10.1093/gbe/evaf125.

O’Donnell, Samuel, Gabriela Rezende, Jean-Philippe Vernadet, Alodie Snirc, and Jeanne Ropars. 2025b. “Harbouring Starships: The Accumulation of Large Horizontal Gene Transfers in Domesticated and Pathogenic Fungi.” Preprint, bioRxiv, January 30. 10.1101/2024.07.03.601904.

O’Donnell, Samuel, Jia-Xing Yue, Omar Abou Saada, et al. 2023. “Telomere-to-Telomere Assemblies of 142 Strains Characterize the Genome Structural Landscape in Saccharomyces Cerevisiae.” Nature Genetics 55 (8): 1390–99. 10.1038/s41588-023-01459-y.

Palmer, Jonathan M., and Jason Stajich. 2020. Funannotate v1.8.1: Eukaryotic Genome Annotation. V. v1.8.1. Zenodo, released September 28. 10.5281/zenodo.4054262.

Park, Chan Ho, Gautam Shirsekar, Maria Bellizzi, et al. 2016. “The E3 Ligase APIP10 Connects the Effector AvrPiz-t to the NLR Receptor Piz-t in Rice.” PLOS Pathogens 12 (3): e1005529. 10.1371/journal.ppat.1005529.

Park, Chan-Ho, Songbiao Chen, Gautam Shirsekar, et al. 2012. “The Magnaporthe Oryzae Effector AvrPiz-t Targets the RING E3 Ubiquitin Ligase APIP6 to Suppress Pathogen-Associated Molecular Pattern–Triggered Immunity in Rice.” The Plant Cell 24 (11): 11. 10.1105/tpc.112.105429.

Pedersen, Thomas Lin. 2025. Ggraph: An Implementation of Grammar of Graphics for Graphs and Networks. https://github.com/thomasp85/ggraph.

Peña, Mariana Villalba de la, Pauliina A. M. Summanen, Martta Liukkonen, and Ilkka Kronholm. 2023. “Chromatin Structure Influences Rate and Spectrum of Spontaneous Mutations in Neurospora Crassa.” Genome Research 33 (4): 4. 10.1101/gr.276992.122.

Peng, Zhao, Ely Oliveira-Garcia, Guifang Lin, et al. 2019. “Effector Gene Reshuffling Involves Dispensable Mini-Chromosomes in the Wheat Blast Fungus.” PLoS Genetics 15 (9): 9. 10.1371/journal.pgen.1008272.

Quinlan, Aaron R., and Ira M. Hall. 2010. “BEDTools: A Flexible Suite of Utilities for Comparing Genomic Features.” Bioinformatics 26 (6): 6. 10.1093/bioinformatics/btq033.

Raffaele, Sylvain, Rhys A. Farrer, Liliana M. Cano, et al. 2010a. “Genome Evolution Following Host Jumps in the Irish Potato Famine Pathogen Lineage.” Science 330 (6010): 1540–43. 10.1126/science.1193070.

Raffaele, Sylvain, Rhys A. Farrer, Liliana M. Cano, et al. 2010b. “Genome Evolution Following Host Jumps in the Irish Potato Famine Pathogen Lineage.” Report. Science 330 (6010): 6010. 10.1126/science.1193070.

Rahnama, Mostafa, Bradford Condon, João P. Ascari, et al. 2023. “Recent Co-Evolution of Two Pandemic Plant Diseases in a Multi-Hybrid Swarm.” Nature Ecology & Evolution 7 (12): 2055–66. 10.1038/s41559-023-02237-z.

Romeijn, Josje, Iñigo Bañales, and Michael F. Seidl. 2025. “Extensive Horizontal Transfer of Transposable Elements Shapes Fungal Mobilomes.” Current Biology 0 (0). 10.1016/j.cub.2025.11.012.

Rowe, David, Jun Huang, Wei Zhang, et al. 2023. “Natural Genomic Variation in Rice Blast Genomes Is Associated with Specific Heterochromatin Modifications.” Preprint, bioRxiv, September 1. 10.1101/2023.08.30.555587.

SAMtoBAM. 2025. SAMtoBAM/Stargraph: V1.0.0-Beta. Zenodo, released June 7. 10.5281/zenodo.15616300.

Sánchez-Vallet, Andrea, Simone Fouché, Isabelle Fudal, et al. 2018. “The Genome Biology of Effector Gene Evolution in Filamentous Plant Pathogens.” Annual Review of Phytopathology 56 (1): 1. 10.1146/annurev-phyto-080516-035303.

Sato, Yukiyo, Roos Bex, Grardy C. M. van den Berg, et al. 2025. “Starship Giant Transposons Dominate Plastic Genomic Regions in a Fungal Plant Pathogen and Drive Virulence Evolution.” Nature Communications 16 (1): 6806. 10.1038/s41467-025-61986-6.

Schloissnig, Siegfried, Samarendra Pani, Jana Ebler, et al. 2025. “Structural Variation in 1,019 Diverse Humans Based on Long-Read Sequencing.” Nature 644 (8076): 442–52. 10.1038/s41586-025-09290-7.

Schotanus, Klaas, Jessica L. Soyer, Lanelle R. Connolly, et al. 2015. “Histone Modifications Rather than the Novel Regional Centromeres of Zymoseptoria Tritici Distinguish Core and Accessory Chromosomes.” Epigenetics & Chromatin 8 (1): 1. 10.1186/s13072-015-0033-5.

Smit, AFA, R Hubley, and P Green. 2015. “RepeatMasker Open-4.0. 2013–2015.” Seattle, USA.

Soyer, Jessica L., Mennat El Ghalid, Nicolas Glaser, et al. 2014. “Epigenetic Control of Effector Gene Expression in the Plant Pathogenic Fungus Leptosphaeria Maculans.” PLoS Genetics 10 (3): e1004227. 10.1371/journal.pgen.1004227.

Sweigard, J A, A M Carroll, S Kang, L Farrall, F G Chumley, and B Valent. 1995. “Identification, Cloning, and Characterization of PWL2, a Gene for Host Species Specificity in the Rice Blast Fungus.” The Plant Cell 7 (8): 1221–33. 10.1105/tpc.7.8.1221.

Torres, David E., Ursula Oggenfuss, Daniel Croll, and Michael F. Seidl. 2020. “Genome Evolution in Fungal Plant Pathogens: Looking beyond the Two-Speed Genome Model.” Fungal Biology Reviews 34 (3): 136–43. 10.1016/j.fbr.2020.07.001.

Urquhart, Andrew S., Nicholas F. Chong, Yongqing Yang, and Alexander Idnurm. 2022. “A Large Transposable Element Mediates Metal Resistance in the Fungus Paecilomyces Variotii.” Current Biology 32 (5): 5. 10.1016/j.cub.2021.12.048.

Urquhart, Andrew S., Emile Gluck-Thaler, and Aaron A. Vogan. 2024. “Gene Acquisition by Giant Transposons Primes Eukaryotes for Rapid Evolution via Horizontal Gene Transfer.” Science Advances 10 (49): eadp8738. 10.1126/sciadv.adp8738.

Urquhart, Andrew S., Samuel O’Donnell, Emile Gluck-Thaler, and Aaron A. Vogan. 2025. “A Natural Mechanism of Eukaryotic Horizontal Gene Transfer.” Preprint, bioRxiv, March 6. 10.1101/2025.02.28.640899.

Urquhart, Andrew S., Aaron A. Vogan, Donald M. Gardiner, and Alexander Idnurm. 2023. “Starships Are Active Eukaryotic Transposable Elements Mobilized by a New Family of Tyrosine Recombinases.” Proceedings of the National Academy of Sciences 120 (15): 15. 10.1073/pnas.2214521120.

Urquhart, Andrew, Aaron A. Vogan, and Emile Gluck-Thaler. 2024. “*Starships*: A New Frontier for Fungal Biology.” Trends in Genetics 40 (12): 1060–73. 10.1016/j.tig.2024.08.006.

Valent, Barbara. 2021. “The Impact of Blast Disease: Past, Present, and Future.” In Magnaporthe Oryzae: Methods and Protocols, edited by Stefan Jacob. Springer US. 10.1007/978-1-0716-1613-0_1.

Vogan, Aaron A., S. Lorena Ament-Velásquez, Eric Bastiaans, et al. 2021. “The Enterprise, a Massive Transposon Carrying Spok Meiotic Drive Genes.” Genome Research 31 (5): 789–98. 10.1101/gr.267609.120.

Wang, Liyuan, Han Chen, JiangJiang Li, et al. 2020. “Effector Gene Silencing Mediated by Histone Methylation Underpins Host Adaptation in an Oomycete Plant Pathogen.” Nucleic Acids Research 48 (4): 1790–99. 10.1093/nar/gkz1160.

Wang, Qinhu, Cong Jiang, Chenfang Wang, Changjun Chen, Jin-Rong Xu, and Huiquan Liu. 2017. “Characterization of the Two-Speed Subgenomes of Fusarium Graminearum Reveals the Fast-Speed Subgenome Specialized for Adaption and Infection.” Frontiers in Plant Science 8 (February). 10.3389/fpls.2017.00140.

Wu, Thomas D., and Colin K. Watanabe. 2005. “GMAP: A Genomic Mapping and Alignment Program for mRNA and EST Sequences.” Bioinformatics 21 (9): 1859–75. 10.1093/bioinformatics/bti310.

Wyk, Stephanie van, Christopher H. Harrison, Brenda D. Wingfield, Lieschen De Vos, Nicolaas A. van der Merwe, and Emma T. Steenkamp. 2019. “The RIPper, a Web-Based Tool for Genome-Wide Quantification of Repeat-Induced Point (RIP) Mutations.” PeerJ 7 (August): e7447. 10.7717/peerj.7447.

Xu, Mengting, Ziyue Sun, Huanbin Shi, et al. 2024. “Two H3K36 Methyltransferases Differentially Associate with Transcriptional Activity and Enrichment of Facultative Heterochromatin in Rice Blast Fungus.” aBIOTECH 5 (1): 1–16. 10.1007/s42994-023-00127-3.

Yang, Li-Chao, Yong-Liang Gan, Li-Yan Yang, Bo-Le Jiang, and Ji-Liang Tang. 2018. “Peptidoglycan Hydrolysis Mediated by the Amidase AmiC and Its LytM Activator NlpD Is Critical for Cell Separation and Virulence in the Phytopathogen Xanthomonas Campestris.” Molecular Plant Pathology 19 (7): 1705–18. 10.1111/mpp.12653.

Yoshida, Kentaro, Hiromasa Saitoh, Shizuko Fujisawa, et al. 2009. “Association Genetics Reveals Three Novel Avirulence Genes from the Rice Blast Fungal Pathogen Magnaporthe Oryzae.” The Plant Cell 21 (5): 1573–91. 10.1105/tpc.109.066324.

Yoshida, Kentaro, Diane G. O. Saunders, Chikako Mitsuoka, et al. 2016. “Host Specialization of the Blast Fungus Magnaporthe Oryzae Is Associated with Dynamic Gain and Loss of Genes Linked to Transposable Elements.” BMC Genomics 17 (May): 370. 10.1186/s12864-016-2690-6.

Zhang, Wei, Jun Huang, and David E. Cook. 2021. “Histone Modification Dynamics at H3K27 Are Associated with Altered Transcription of in Planta Induced Genes in Magnaporthe Oryzae.” PLOS Genetics 17 (2): 2. 10.1371/journal.pgen.1009376.

Zhang, Yong, Tao Liu, Clifford A. Meyer, et al. 2008. “Model-Based Analysis of ChIP-Seq (MACS).” Genome Biology 9 (9): 9. 10.1186/gb-2008-9-9-r137.

Zhong, Zhenhui, Meilian Chen, Lianyu Lin, et al. 2018. “Population Genomic Analysis of the Rice Blast Fungus Reveals Specific Events Associated with Expansion of Three Main Clades.” The ISME Journal 12 (8): 8. 10.1038/s41396-018-0100-6.

Zong, Xifang, Yaxin Lou, Mengshuang Xia, et al. 2024. “Recombination and Repeat-Induced Point Mutation Landscapes Reveal Trade-Offs between the Sexual and Asexual Cycles of *Magnaporthe Oryzae*.” Journal of Genetics and Genomics 51 (7): 723–34. 10.1016/j.jgg.2024.03.003.

